# Dynamics of genotype frequencies in finite partially clonal populations under mutation and genetic drift

**DOI:** 10.64898/2026.04.10.717696

**Authors:** Solenn Stoeckel, Jean-Pierre Masson

## Abstract

Most eukaryotes reproduce using partial clonality. Yet we still lack population genetic models capable of efficiently reconstructing and predicting genotype frequency changes across generations, and of elucidating underlying evolutionary processes in longitudinally monitored partially clonal populations. Here, we develop a Wright-Fisher-like model describing the mean and variance of genotype frequencies through time at a single multi-allelic locus in finite partially clonal populations with mutation.

We show that different initial conditions and clonality rates generate distinct genotype-frequency trajectories. Regardless of clonality rate, partially clonal populations converge toward the same equilibrium as fully sexual populations, determined solely by reciprocal mutation rates among alleles. Dynamics proceed in two phases: genotype frequencies first approach Hardy-Weinberg proportions, then evolve along the Hardy-Weinberg parabola until reaching equilibrium. Mean allele frequencies and gene diversity remain unaffected by clonality throughout these trajectories. Instead, clonality rate determines the speed at which populations return to Hardy-Weinberg proportions, thereby shaping the sequence of genotype-frequency distributions traversed over successive generations.

The variance around mean trajectories depends on parental genotype frequencies and population size, and is not directly determined by clonality rate; rather, clonality affects it only indirectly by altering the sequence of genotype distributions traversed. This explains why partially clonal populations may exhibit both positive and negative F_IS_ at different loci, and why variance in F_IS_ among loci provides a robust proxy for inferring clonality rates.

Our model provides a theoretical framework for analysing, predicting and interpreting temporal changes in genotype frequencies in monitored populations, establishing a foundation for future inference methods based on time-series genotyping data.

## 1 Introduction

Population monitoring, i.e., the repeated survey of a given population over time using standardized methods, is fundamental in ecology and evolution (Murray and Krebs 2024). It enables investigation of demography, life-history evolution, responses to environmental change, and phenotypic and genetic changes across generations, and, in applied contexts, provides a basis for evaluating management actions targeting the focal population. Beyond the focal population and species, monitoring programs can yield insights into ongoing ecological interactions and offer empirical evidence of environmental change (Moussy et al. 2022). As genotyping technologies become increasingly automated and affordable, many population monitoring programs now incorporate temporal genetic data at multiple markers (e.g., example of past studies: Becheler et al. 2017; Pearman et al 2024; Saubin et al. 2024; Bocharova et al. 2025; Shainker-Connelly et al. 2025). This trend underscores the need for population genetic models capable of describing the temporal dynamics of genetic diversity across generations while accounting for species-specific biological characteristics.

The mode by which individuals in a population transmit their genetic information across generations, i.e. reproductive modes, determines the evolution of genetic diversity and its structure within species, and constrains the adaptive potential of species facing environmental change (Hamrick and Godt 1996; Duminil et al. 2007; Eckert et al. 2010; Ellegren and Galtier 2016; Orive et al. 2017). Understanding and predicting the impacts of reproductive modes on genetic diversity is therefore central to ecology and evolution (Maynard Smith 1978; Otto and Lenormand 2002; Diaz-Martin et al. 2023; Chaparro-Pedraza et al. 2024).

Clonality is a reproductive mode in which offspring are generated by mitosis and are genetically identical to their single parent, except for rare mitotic mutations and recombination events (Richards 2003; DeMeeus et al. 2007; Stoeckel et al. 2021a). Most eukaryotes, including many primary producers, ecosystem engineers, pathogens and invasive species, reproduce using partial clonality, producing clonal and sexual offspring either sequentially or simultaneously within their life cycle (Avise 2008; Schön et al. 2009; Tibayrenc et al. 2015; Orive & Krueger-Hadfield 2021). In such populations, reproduction is characterized by a rate of clonality, defined as the proportion of clonal offspring among all offspring produced by individuals capable of both clonal and sexual reproduction (Marshall & Weir 1979, Stoeckel et al. 2021a). Because clonality directly affects genotype transmission, the evolution of genetic diversity, both forward and backward in time, requires explicit tracking of population genetic states defined by genotype frequencies, rather than allele-based approximations, that are sufficient for fully sexual populations (Ewens 2004; Stoeckel & Masson 2014). Interestingly, the genetic evolution of partially clonal populations cannot be described as a simple linear combination of the dynamics expected under fully sexual and fully clonal reproduction (Marshall and Weir 1979; Orive 1993; Berg and Lascoux 2000; Balloux et al. 2003; Bengtson 2003; Ceplitis 2003; Stoeckel and Masson 2014; Hartfield et al. 2016; Abdalrahem et al. 2025).

At each generation, genotype frequencies resulting from sexual and clonal reproduction are combined within a common offspring pool, generating evolutionary dynamics and genotype-frequency trajectories determined by rate of clonality. The exact distributions of genotype frequencies across generations and at equilibrium can be obtained from a Markov chain model (Stoeckel and Masson 2014). However, the number of possible genetic states of the population increases rapidly with population size *N* and the number of alleles *k*:

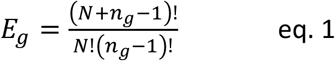

with *n*_*g*_, the number of possible genotypes, being equal to 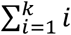 for diploids.

With only two alleles, for *N* = 2, *E*_*g*_ = 6; for *N* = 3, *E*_*g*_ = 10; for *N* = 100, *E*_*g*_ = 5151;

And with four alleles, for *N* = 2, *E*_*g*_ = 55; for *N* = 3, *E*_*g*_ = 220; for *N* = 100, *E*_*g*_ > 4.26 × 10^12^; which makes the calculations and computations complicated or even impossible (Reichel et al. 2015). Rather than analyzing the probabilities associated with discrete genotypic states, it is more appropriate to work with continuous genotype frequency distributions and their associated probabilities, as previously developed for panmictic sexual populations (e.g., Ewens 2004; Fu et al. 2003) and partially selfing populations (Pollak 1987).

In this study, building on our previous discrete framework of population genetic equations for partially clonal populations at a single locus (Stoeckel and Masson 2014), we extend the model to a continuous representation of genotype frequency distributions at a multi-allelic locus, an approach better suited for large population sizes. We characterize the temporal evolution of mean genotype frequencies and their dispersion across different rates of clonality (from exclusively sexual to exclusively clonal populations), representing this dispersion as concentration ellipses. We then focus on the analytically tractable case of a diallelic locus, for which we derive analytical expressions and examine in detail how classical population genetic summary statistics evolve under partial clonality. From these results, we identify key model properties with potential to inform future inference methods for empirical datasets from populations repeatedly genotyped across successive generations. Finally, because partial clonality requires explicit tracking of genotype (rather than allele) frequencies, we complement our framework with graphical representations of mean and variance dynamics at the genotype scale to aid interpretation, again for the two-allele case.

## 2. Model

For this model, we consider only genetic diversity that is neutral in the sense of temporally conditional neutrality (De Lafontaine et al. 2018), i.e., genetic variants conferring no selective advantage or disadvantage to any individual over the spatial and temporal scales across which the population is monitored.

### 2.1. The genetic system and its model for one locus with multiple alleles

First, we consider a single locus with a fixed number of alleles *k* through discrete (non-overlapping) generations in a finite diploid population without selection, reproducing at clonality rate *c*. This framework extends the classical Wright-Fisher model, first formulated by Fisher (1922) and formalized by Wright (1931), expanded here to include multiple alleles and partial clonality. Since clonality implies that genotypes rather than alleles are transmitted across generations, our equations are formalized considering genotypes and not alleles, as more commonly considered in classical Wright-Fisher model (Ewans 2004; Stoeckel and Masson 2014).

Specifically, we consider *k* ≥ 2 alleles, denoted *A*_1_, ⋯, *A*_*i*_, ⋯ *A*_*k*_ with equal and symmetric mutation rate between alleles. All alleles mutate at a fixed rate *u*_*ij*_ = *u*, for every pair (*A*_*i*_ → *A*_*j*_, *A*_*j*_ → *A*_*i*_) with *i* ≠ *j*, and remain unchanged with rate *u*_*ii*_ = 1 − (*k* − 1)*u* = *v*. It corresponds to a K-allele mutation (KAM) model (Putman and Carbone, 2014; Weir and Cockerham, 1984), which accurately describes mutation patterns at both microsatellite and SNP loci. In such mutation model, mutation rates can be interpreted as approximating the “*disturbing factor of gene [allele] frequencies*” in the sense of Wright (1931), potentially subsuming limited immigration from external populations. To facilitate the analysis of the underlying processes, we assume throughout that the reproductive population size *N* remains constant across generations. However, our framework can be readily extended to accommodate time-varying population sizes, i.e., *N*^(*n*)^ as required for realistic applications to monitored populations. By contrast, incorporating such variation into a Markov chain framework would be computationally intensive, as it would require rectangular transition matrices to be redefined at each generation.

We define the collection 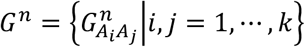 of random variables 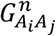 representing the number of individuals of genotype *A*_*i*_*A*_*j*_ at generation *n*. This set includes:

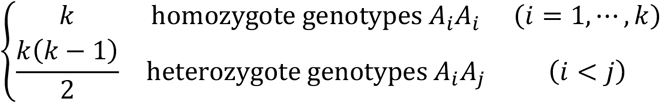

with 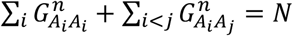, the finite population size.

Population geneticists and ecologists are primarily concerned with genotype frequencies 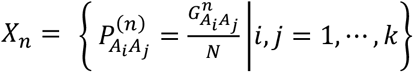, of types 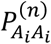 and 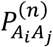 representing the proportions of homozygotes and heterozygotes, respectively, at generation *n* ∈ ℕ (with *n* ≥ 0) in the population under study, with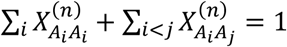 .

If *X*_*n*_ takes the values 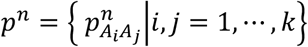 the expected genotype frequencies of homozygotes and heterozygotes in the pool of potential offspring at generation *n* + 1 are given by the following system of equations:

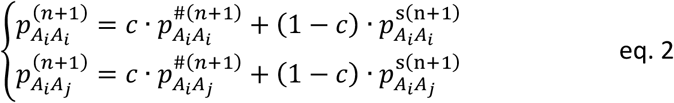

where 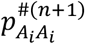 and 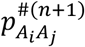 are the genotype frequencies among potential offspring produced by clonal reproduction in the next generation:

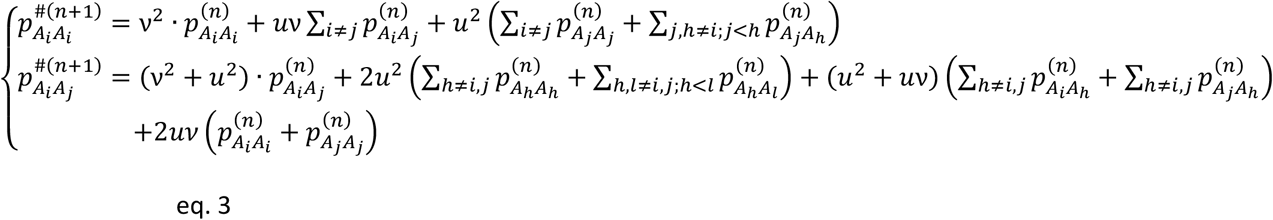

and where 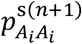and 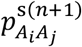 are the genotype frequencies among potential offspring produced by sexual reproduction:

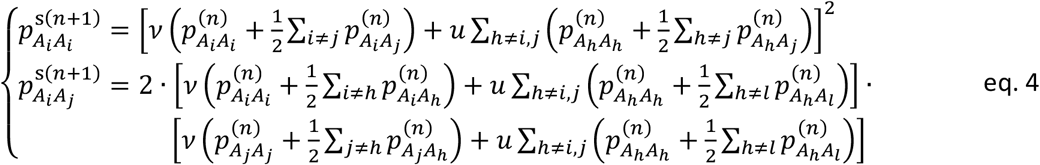

which simplify to:

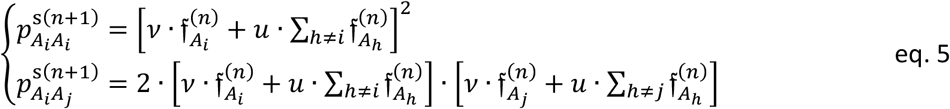

where 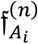 denotes the frequency of the *i*^*th*^ allele at generation *n*, defined as:

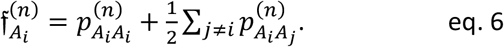

The sequence (*X*^*n*^)_*n*≥0_ is a Markov chain taking values in the simplex 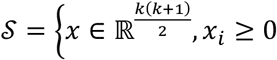 for 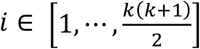 and 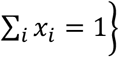 .

Under the assumption that individuals are sampled independently, *G*^*n*+1^ follows a multinomial distribution *ℳ*(*N, p*^*n*+1^; *c*; *u*) with *E*(*G*^*n*+1^) = *N* · *p*^*n*+1^. The *G*^*n*+1^ variance covariance matrix has diagonal elements 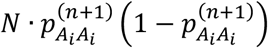 with *i* = 1, ⋯, *k* and off-diagonal elements 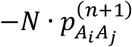 with *i, j* = 1, ⋯, *k. X*^*n*^ is fully determined by the initial condition *p*^0^.

### 2.2. The tractable case of one locus with two alleles

To investigate the dynamics of the system and accurately represent the trajectories of genotype frequencies, for the remainder of this study, we focus on the tractable case of a single locus with two alleles. We thus define 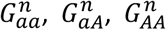 as the stochastic variables accounting for the number of individuals with genotypes aa, aA and AA at generation *n*, with 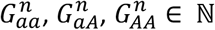 and 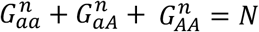 . We simplify notations by denoting 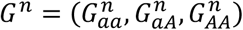 the stochastic variables representing the count aa, aA and AA individuals within the population at generation *n*, and the corresponding genotypic frequencies as 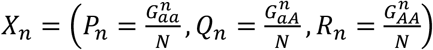 .

If at generation *n, X*_*n*_ takes the values (*p*_*n*_, *q*_*n*_, *r*_*n*_), hereafter referred to as a distribution of genotype frequencies, then the genotype frequency distribution at generation *n* + 1, (*p*_*n*+1_, *q*_*n*+1_, *r*_*n*+1_) following mutation is given by:

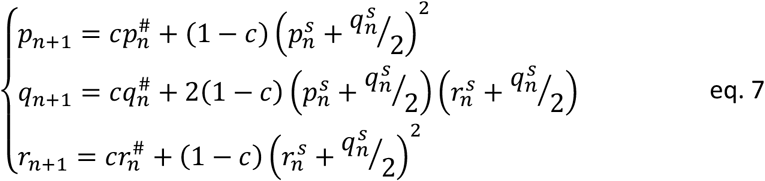

where 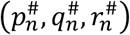 are the genotype frequencies among potential offspring produced by clonal reproduction:

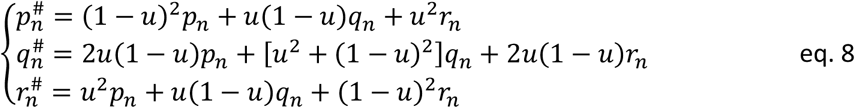

and where(*p*^s^, *q*^s^, *r*^s^) are the genotype frequencies among potential offspring produced by sexual reproduction:

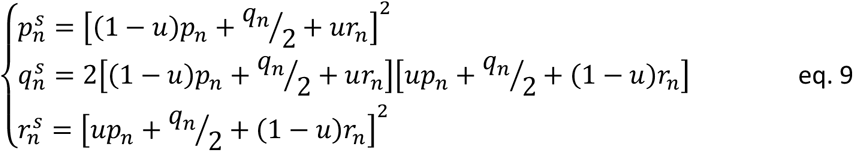

For a given initial conditions (*p*_0_, *q*_0_, *r*_0_), a rate of clonality *c* and a mutation rate *u*, (*p*_*n*+1_, *q*_*n*+1_, *r*_*n*+1_) depends only on (*p*_*n*_, *q*_*n*_, *r*_*n*_), and (*X*_*n*_)_*n*≥0_ is a Markov chain taking values in the simplex *S* = {*x* ∈ ℝ^3^, *x*_*i*_ ≥ 0 and *x*_1_ + *x*_2_ + *x*_3_ = 1}.

Under the assumption that individuals are sampled independently, *G*^*n*+1^ follows a multinomial distribution *ℳ* (*N*, (*p*_*n*+1_, *q*_*n*+1_, *r*_*n*+1_); *c, u*). Therefore, the probability of the event *G*^*n*+1^ = (*s*_*aa*_, *s*_*aA*_, *s*_*AA*_) at generation *n* + 1 given that the parental genotype counts at generation *n* were *G*^*n*^ = (*r*_*aa*_, *r*_*aA*_, *r*_*AA*_) is:

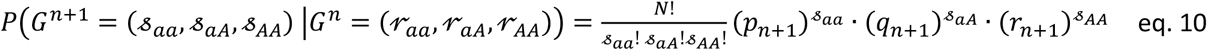

with *s*_*aa*_ + *s*_*aA*_ + *s*_*AA*_ = *N, r*_*aa*_ + *r*_*aA*_ + *r*_*AA*_ = *N* and *p*_*n*+1_ + *q*_*n*+1_ + *r*_*n*+1_ = 1. The mean and variance-covariance matrix are, respectively,

- 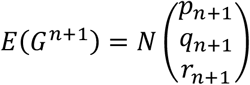
- 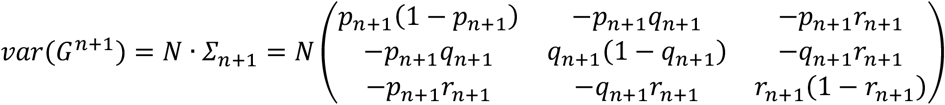

The multinomial distribution *ℳ*(*N*, (*p*_*n*_, *q*_*n*_, *r*_*n*_); *c, u*) is well approximated by a multinormal distribution *Y*^*n*^ ∝ *N* (*Nμ, NΣ*) with 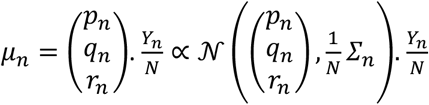 takes it values within the two-dimensional simplex; *rg*(*Σ*_*n*_) = 2.

The sequence (*p*_*n*+1_, *q*_*n*+1_, *r*_*n*+1_) = *F*(*p*_*n*_, *q*_*n*_, *r*_*n*_; *c, u*) defined by equation 7 is bijective and globally invertible. When *n* → ∞, it converges to the unique stable fixed point 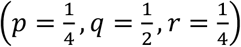 . In the following proof, we denote this stable fixed point as 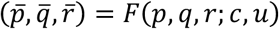 to simplify the notation.

Setting that *r* = 1 − *p* − *q*, the recurrence then becomes 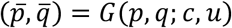 :

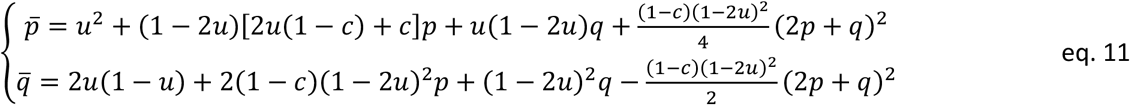

In the following, we will assume that 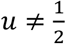 .

A straightforward computation from this recurrence gives:

- the transition of the allele frequency from one generation to the next:

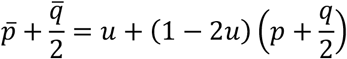

as well as its inverse:

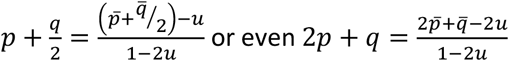
- Moreover, since the sequence is arithmetico–geometric:

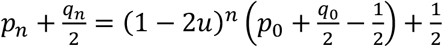

So when *n* → +∞, the allele frequency tends to 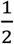

We therefore obtain the inverse of *G* explicitely:

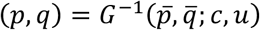

By replacing 2*p* + *q* with 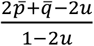 in the system of equations 11 resulting in:

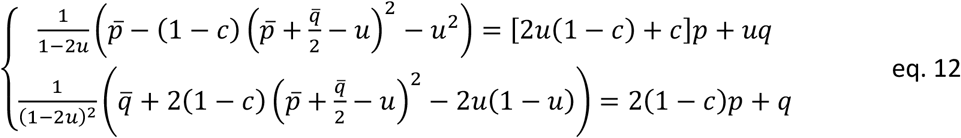

Equation 12 is a system of two linear equations in the unknowns *p* and *q*. Its solution explicitly gives *p* and *q* as function of 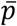 and 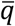, and the parameters *c* and *u*.

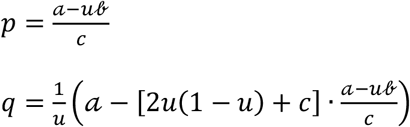

With 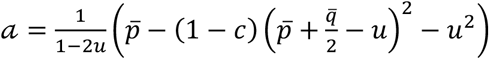

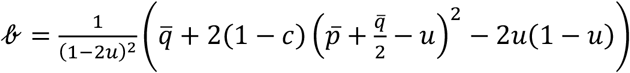

Therefore, the map *G* is a bijection on Δ = {(*p, q*) ∈ ℝ^2^_≥0_ : *p* + *q* ≤ 1}, and thus *F* is also bijective on the simplex *S* = {(*p, q, r*) ∈ ℝ^3^_≥0_ : *p* + *q* + *r* = 1}

To find fixed points, we set 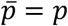 and 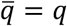 in 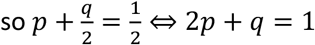 and the solution of (12) with 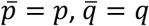 and 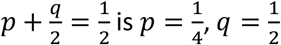 .

It follows that there is a unique fixed point within the simplex *S*, which is 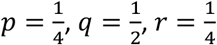 .

The functions *F* (respectively *F*^−1^) and *G* (respectively *G*^−1^) are continuous.

The sequence 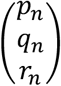 is bounded (it lies in the triangle ABC) and has a unique fixed point 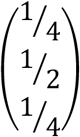, which is the unique accumulation point. Therefore, the sequence 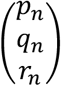 is convergent and converges to 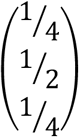 .

Moreover, the determinant of the Jacobian matrix *J* of *G* is equal to:

*det J* = *c*(1 − 2*u*)^3^ at every point of Δ.

For all *c* ≠ 0 and 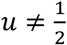, *det J* ≠ 0 therefore *G* and hence *F* are locally invertible, as already shown above.

The fixed point is locally attractive because the eigenvalues of the Jacobian matrix at the fixed point, *λ*_1_ = 1 − 2*u* and *λ*_2_ = *c*(1 − 2*u*)^2^ are less than 1 when *c* ≠ 0.

‖*F*(*p, q, r*; *c, u*)‖ → ∞ when ‖(*p, q, r*)‖ → ∞ and *det J* ≠ 0. Then, by Hadamard’s Lemma, *F* is thus globally inversible for all *c* ∈ (0,1) and 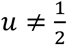 .

### 2.3. Visual representation in two-dimensions

To facilitate the representation and interpretation of our results, we work in two dimensions within the vector subspace associated with the plane of the simplex *S* (Figure 1). (*p, q, r*) are the coordinates of the generic point of the simplex *S* in (*e*_1_, *e*_2_, *e*_3_), the canonical basis of ℝ^3^. For convenience, we introduce a new basis (*e*_1_′, *e*_2_′, *e*_3_′), in which the vectors *e*_*1*_*′, e*_*2*_*′*, and *e*_*3*_*′* are respectively parallel to and pointing in the same direction as 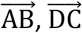, and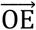 .

**Figure 1.**
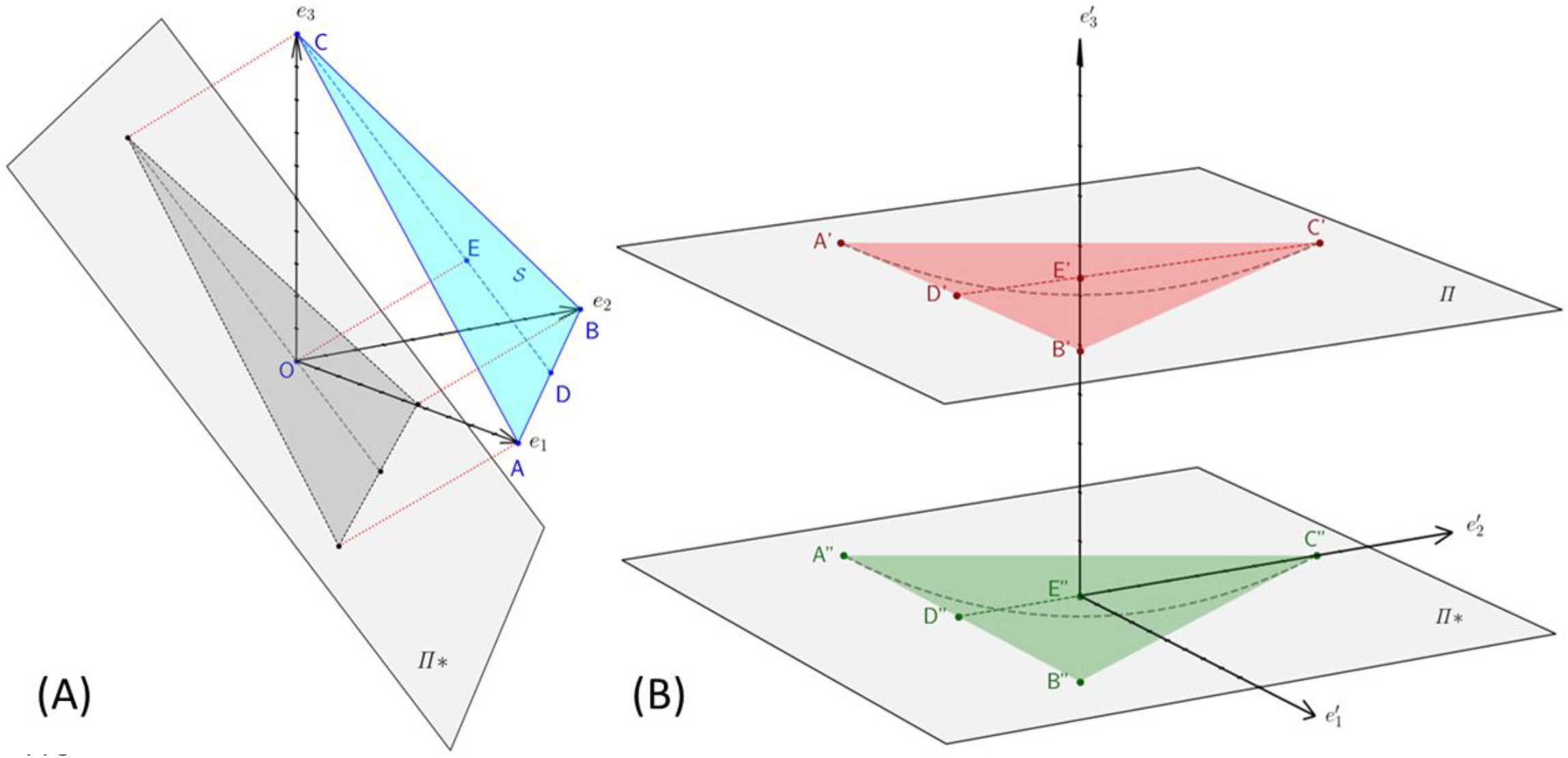
Visual representation of the sub-vector space associated with the plane containing the genotype frequency state space *S*. **(A)**: Projection from the canonical basis (in blue) onto Π^*^ (in grey). Points 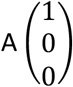, 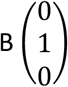 and 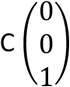 denote the endpoints of the vectors *e*_1_, *e*_2_, *e*_3_. Point 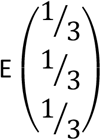, the barycenter of the triangle ABC, projects onto O. **(B)**: Planes *Π* and *Π*^*^, the Hardy-Weinberg parabola is shown as a dashed line in the new basis (*e*_1_′, *e*_2_′, *e*_3_′). *Π*^*^ is spanned by *e*^′^, *e*^′^ ; *Dim Π*^*^ = 2, *e*_3_′ is not required.

It results that

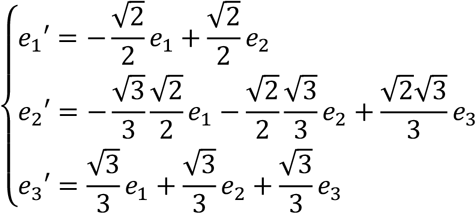

In this new basis 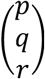has coordinates 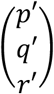 .

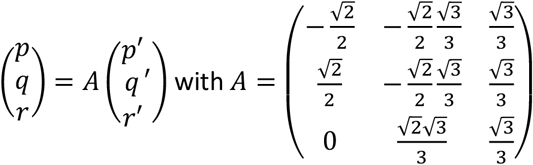

*A* is an orthogonal matrix; so *A*^−1^ = *A*^*t*^ and

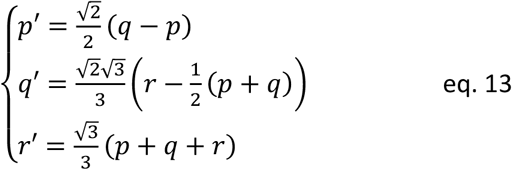

In (*e*_1_ ′, *e*_2_ ′, *e*_3_ ′), *S* belongs to the plane 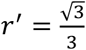 . Π^*^ is spanned by *e*_1_ ′, *e*_2_ ′, associated with the plane 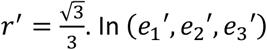. In (*e*_1_ ′, *e*_2_ ′, *e*_3_ ′), the vertices of triangle ABC, delimited space of all possible triplets (*p, q, r*), have for coordinates:

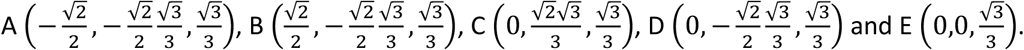

In this new subspace Π^*^, let *p* = π^2^, *q* = 2π(1 − π) and *r* = (1 − π)^2^ be the genotype frequencies expected under Hardy-Weinberg assumptions, namely infinite population size, panmixia and absence of mutation, migration and selection (Hardy 1908; Weinberg 1908), with π being the probability of allele a and 1 − π the probability of allele A.

In (*e*_1_′, *e*_2_′), *p* and *q* become *p*′ and *q*′:

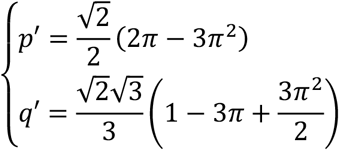

We now perform a translation 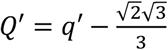 followed by a −30° rotation:

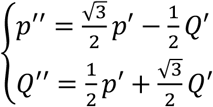

Then:

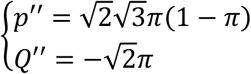

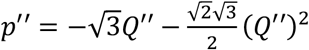 is the Hardy-Weinberg parabola given in our last coordinate system. It passes through the vertex 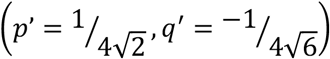 .

The equilateral triangle ABC, the Hardy-Weinberg parabola and all the trajectories are plotted in Figure 2, that summarizes the previous developments.

**Figure 2.**
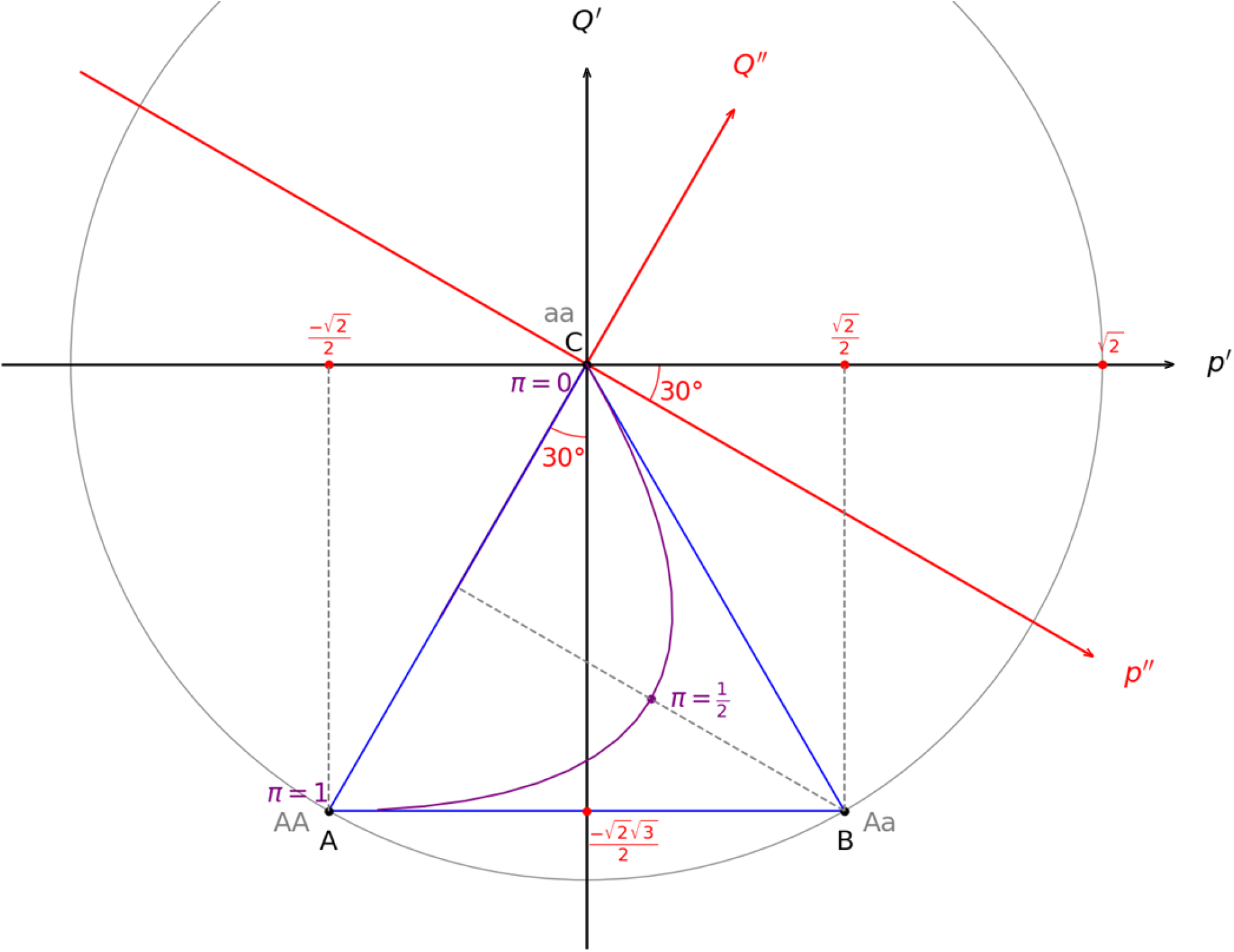
Triangle ABC shown in the coordinate system relative to *(b)* (black axes) and in the coordinate system relative to *(c)* (red axes).

For each 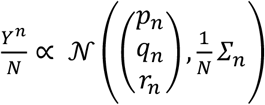, we omit the index n in the remainder of this paragraph as only the location of the mean point is relevant to the following analysis.

Let *λ*_1_ and *λ*_2_ with 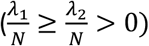 denote the two non-zero eigenvalues of, the variance-covariance matrix *Σ* and let *w*_1_ and *w*_2_ be the corresponding normalized eigenvectors. Both eigenvectors belong to Π^*^. In the coordinate system defined by (*w, w* ), the binormal dispersion distribution 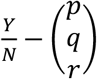 has the following probability density function:

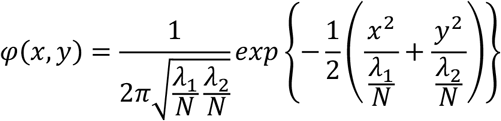

The dispersion around the mean point 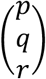 can be visualized and quantified using concentration ellipses, which are homothetic to the reference ellipse defined by:

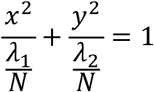

with a positive scaling factor (see Appendix).

The major and minor axes of this reference ellipse are aligned with *w*_1_ and *w*_2_ respectively, and its semi-major and semi-minor axes have lengths 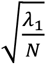, and 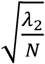, respectively.

The area of the reference ellipse *A* is:

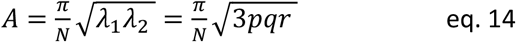

Its eccentricity *e*, bounded between zero and one (where *e* = 0 corresponds to a circle, *e* = 1 to a parabola and *e* > 1 to a hyperbola), measures the deviation of the concentration ellipse from circularity:

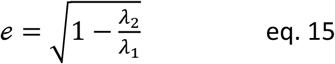

In the following sections, the trajectories of successive (*p*′, *q*′) and their concentration ellipses are displayed in the two-dimensional space Π^*^ (Figure 1).

### 2.4. Trajectories of genotype frequencies in finite mutating populations

We now define properties of trajectories in *S* seen in *Π*^*^. A trajectory is the sequence of genotype frequency distributions across generations within a given population. For one locus with two alleles in diploids, it corresponds to the sequence of (*p*_*n*_, *q*_*n*_, *r*_*n*_) along the unique trajectory leading from the initial point (*p*_0_, *q*_0_, *r*_0_) to its steady-state equilibrium, for a given clonality rate *c* and mutation rate *u* (Figure 3). The mean trajectory and its dispersion, represented by concentration ellipses, can equivalently be displayed in the plane Π^*^ (Figure 4).

**Figure 3.**
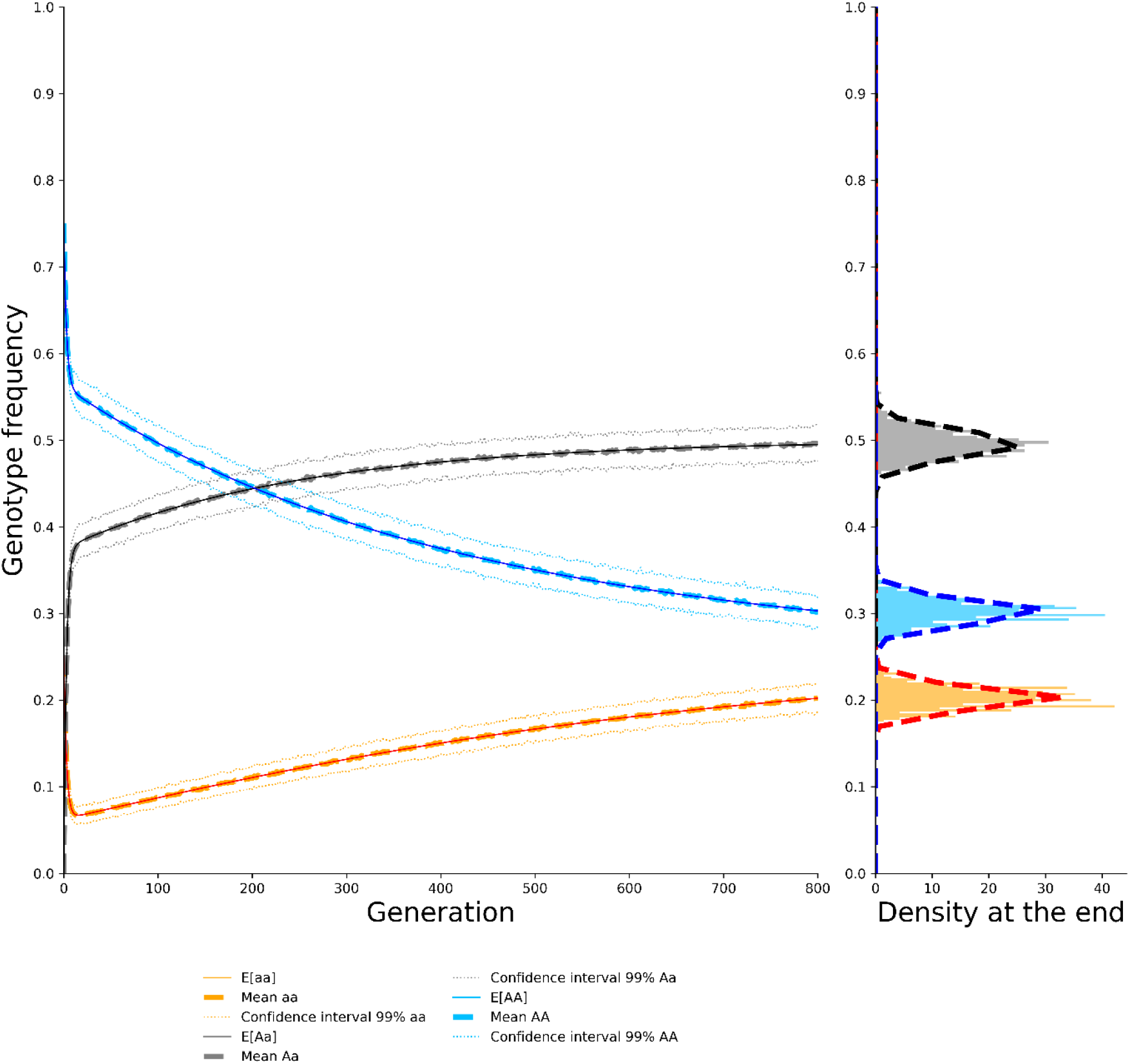
*Left panel*: Temporal dynamics of genotype frequencies across generations (corresponding to Figure 3) for *N* = 1000, *u* = 10^−3^, *c* = 0.7 monitored from generation 0 with initial genotype frequencies (aa=0.25, Aa=0., AA=0.75). Solid lines show the mean genotype frequencies across time (eq.7); Thick dashed lines represent the mean of 300 multinomial draws at each generation. Thin dotted lines show the 99% confidence intervals for each genotype frequency time series, computed from the 300 multinomial draws. Shades of blue correspond to AA, shades of grey to Aa and shades of orange to aa. *Right panel:* Histogram of marginal genotype frequency distributions at generation 800, computed from the 300 multinomial draws (but see Appendix D). Thick dashed lines show the marginal Beta distributions *Beta*(*α*_*k*_, 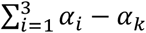 ) of a Dirichlet distribution *Dir*(*α*_0_, *α*_1_, *α*_2_) fitted on these 300 multinomial draws.

**Figure 4.**
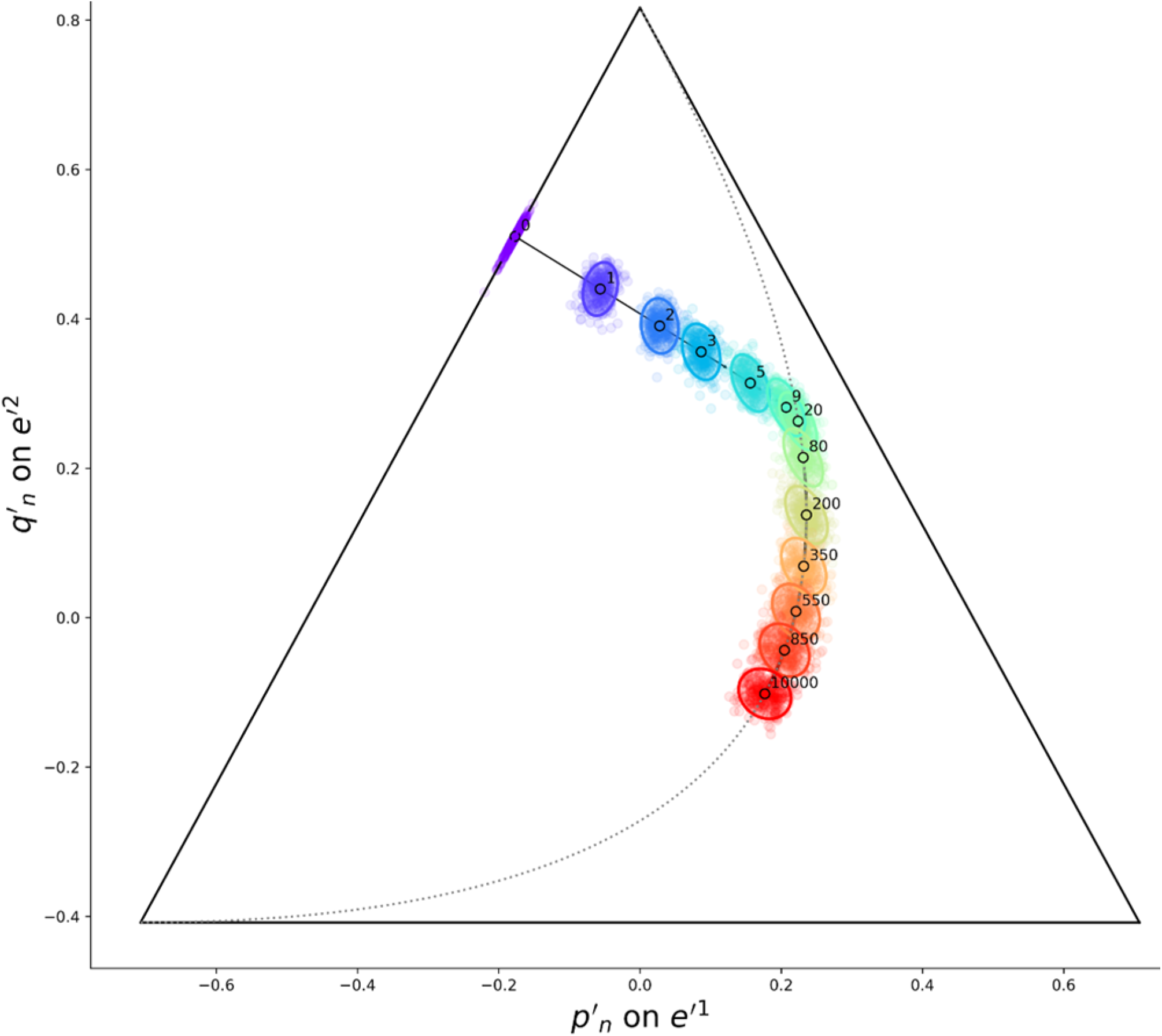
Mean trajectory of genotype frequencies (black line with hollow dots) and associated concentration ellipses across generations (color-coded from violet to red following generation order) in the plane Π^*^ for a population with parameters *N* = 1000, *u* = 0.001, *c* = 0.7, and monitored from generation 0 with initial genotype frequencies (aa=0.25, Aa=0., AA=0.75). For each generation (labeled in black), 300 points sampled from the multinomial model are shown as a scatter plot, illustrating the goodness of fit between multinomial distributions and the multinormal concentration ellipses.

First, we define the direction of a trajectory as the angle *D*, in degree, subtended by the segment connecting two successive genotype frequency distributions in the basis *e*_1_′, *e*_2_′ of *Π*^*^.

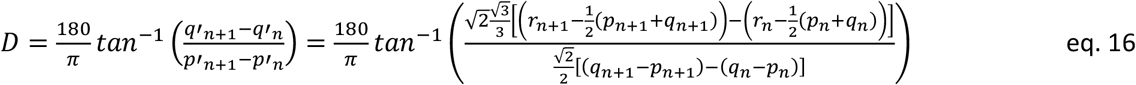

Then, we define the speed of the trajectory as the magnitude of the displacement between two successive generations:

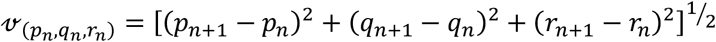

In the subspace *Π*^*^, following the change of orthogonal basis, this speed is given by:

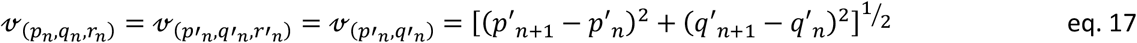

### 2.5. Population genetic indices

To interpret our results in terms of genetic diversity dynamics, we track classical population genetic indices across generations. Specifically, we track *p*_*n*_, the frequency of homozygote *aa; Ho*_*n*_ = *q*_*n*_, the observed heterozygosity; 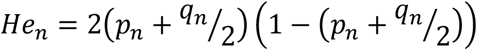 the gene diversity or expected heterozygosity; And Wright’s inbreeding coefficient 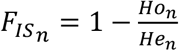 .

We also track the number of generations *n*_*HW*_ required to reach the Hardy-Weinberg parabola as

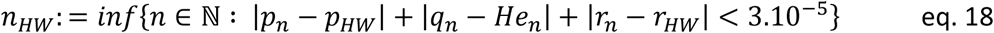

with 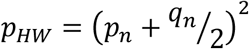 and 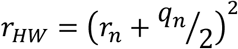, and the number of generations required to reach the stable equilibrium as

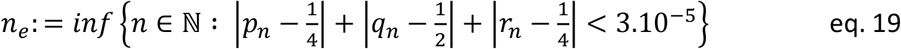

All the equations and computation are implemented in a python library, DyGenClon, which also provides functions for reproducing all figures: https://forge.inrae.fr/solenn.stoeckel/dygenclon

## 3. Results

We present results for a single locus with two alleles, which allows visualization and interpretation through two-dimensional plots. All dynamics and concepts presented for two alleles generalize to a multi-allelic locus (see derivations in section 2.1).

In finite-sized populations under partial clonality, all mean genotype frequency trajectories initiated away from Hardy-Weinberg proportions exhibit two distinct phases: an initial convergence toward Hardy-Weinberg proportions, followed by an evolution constrained to the Hardy-Weinberg parabola (Figure 5). For any given population size, mutation rate and rate of clonality, mean trajectories originating from distinct but closely similar states are unique (bijective). All trajectories diverge from the boundary of the domain to the unique equilibrium point (Figure 6). Trajectories converge only upon reaching the equilibrium point (zooms in Figure 6), enabling both backward and forward tracking of the temporal expectations of genotype frequencies.

**Figure 5:**
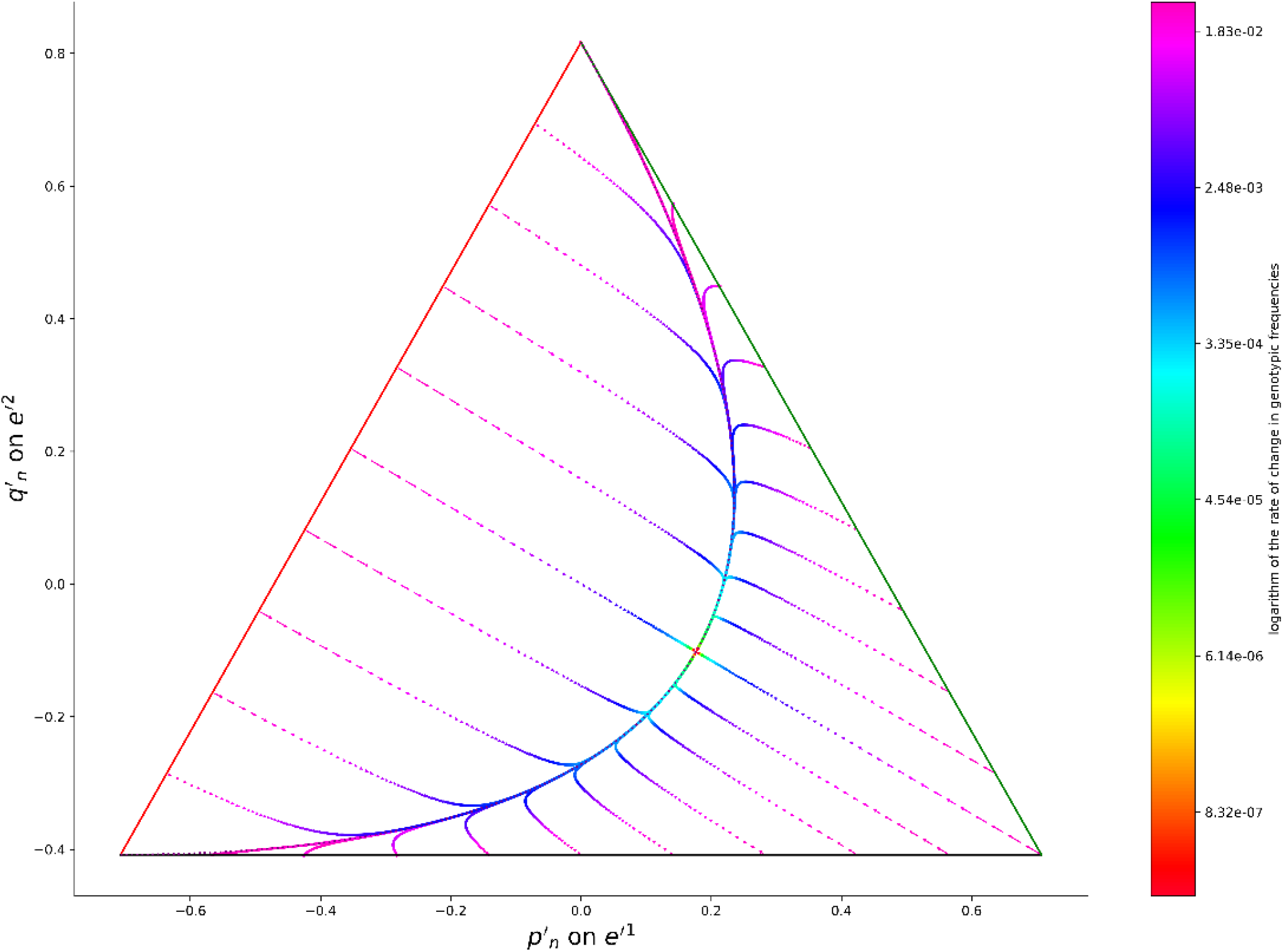
Field of mean trajectories for a population with clonality rate c=0.95 and mutation rate u=10^-3^. Each arrow represents the change in genotype frequencies over one generation. Arrow color indicates the logarithm of the rate of change in genotype frequencies. The Hardy-Weinberg parabola is shown as a dotted grey line. All trajectories converge towards (AA:1/4, Aa:1/2, aa:1/4), corresponding to a zero rate of change in genotype frequencies (red dot), located on the Hardy-Weinberg parabola.

**Figure 6:**
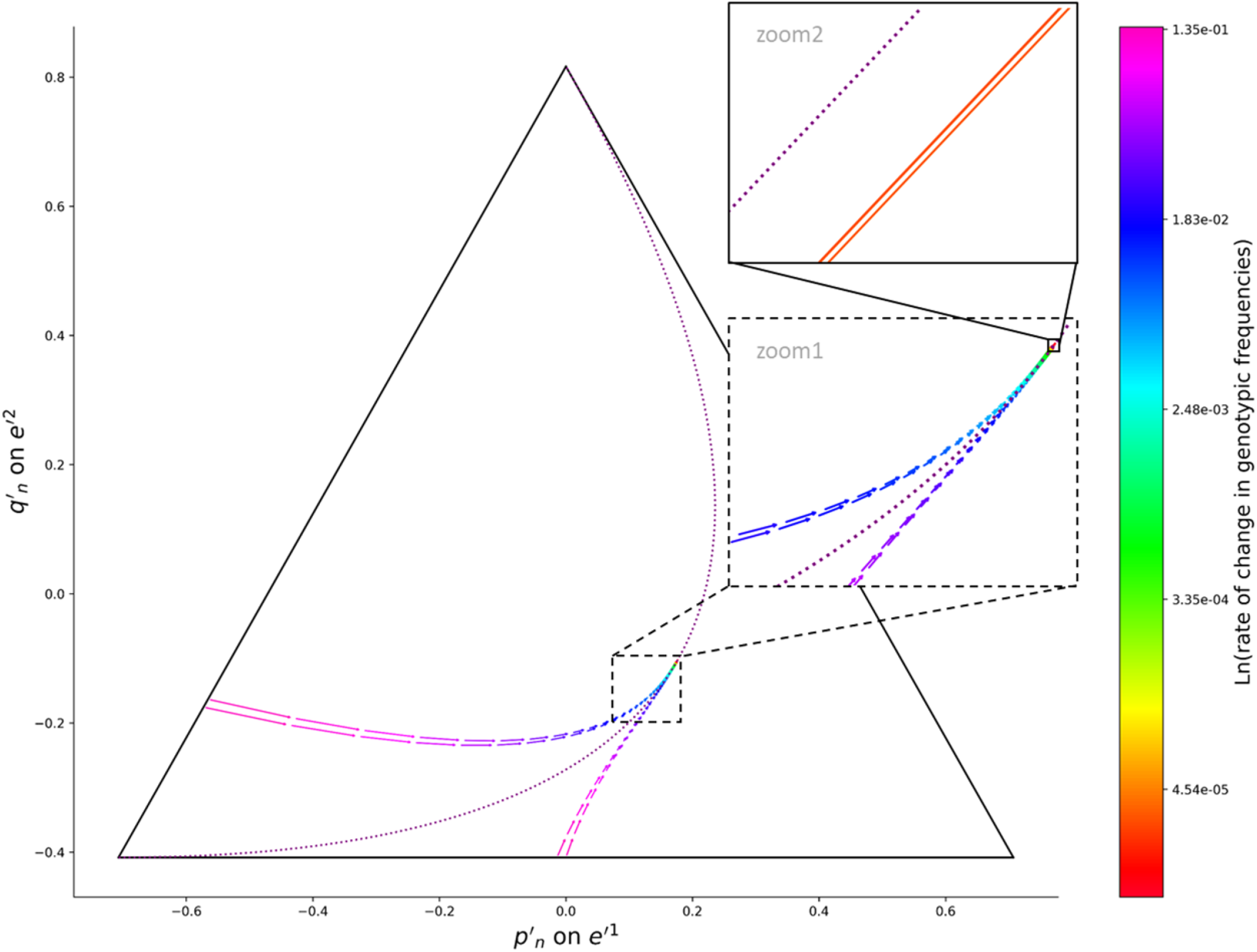
Illustration of the bijective nature of mean trajectories for *c* = 0.95 and *u* = 0.05. Two trajectories originating from closely similar but distinct initial states meet only at the stable equilibrium (AA:1/4, Aa:1/2, aa:1/4). The two successive close-up views reveal that mean trajectories of partially clonal populations do not iterate precisely on, but in the vicinity of the Hardy-Weinberg parabola (dotted purple line, shown centrally in inset 1 and in the top-left of inset 2), never crossing except at the equilibrium point. The first pair of trajectories, to the left of the Hardy-Weinberg parabola, originates at (aa:0.8, Aa:0, AA:0.2) and (aa:0.81, Aa:0, AA:0.19). The second pair of trajectories, to the right of the Hardy-Weinberg parabola, originates at (aa:0.5, Aa:0.5, AA:0) and (aa:0.51, Aa:0.49, AA:0).

A shift in trajectory direction occurs upon reaching the Hardy-Weinberg parabola. The abruptness of this directional change depends on the initial genotype frequencies and the angle at which the trajectory intersects the parabola (Figure 7, Figure S1).

**Figure 7:**
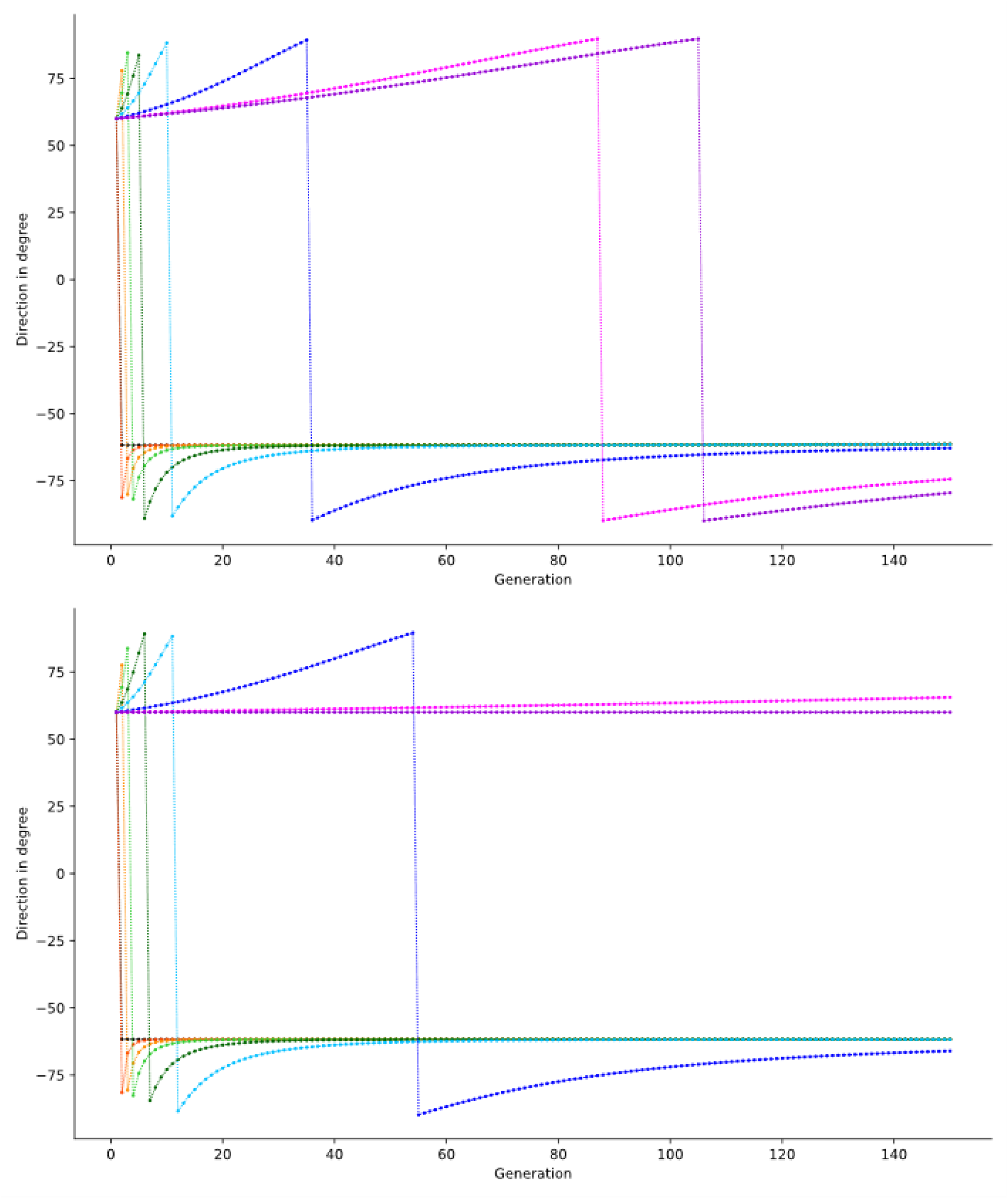
Angular direction of the mean trajectories (*D*, eq.13) originating from the initial point (aa:0.2, Aa:0, AA:0.8). A sudden change in direction occurs upon reaching the Hardy-Weinberg parabola. The top panel corresponds to a mutation rate of 10^−3^ and the bottom panel to a mutation rate of 10^−6^. Line colors represent different clonality rates: black: c=0, orange: c=0.5, gold: c=0.7, limegreen: c=0.8, darkgreen: c=0.9, skyblue: c=0.95, blue: c=0.99, fuchsia: c=0.999, darkviolet: c=1.

In partially and fully clonal populations, for a given population size, mutation rate and rate of clonality, convergence toward the Hardy-Weinberg parabola is rapid when populations are far from it but decelerates as they approach it (Figure 8, Figure S1). At a fixed mutation rate, higher clonality rate increases the mean number of generations required for genotype frequencies to return to the Hardy-Weinberg parabola (Table 1). Increasing mutation rate accelerates the return to the Hardy-Weinberg parabola, as mutation in the K-allele model effectively randomizes allele combinations within individuals carrying mutation, regardless of their mode of reproduction.

**Table 1.**
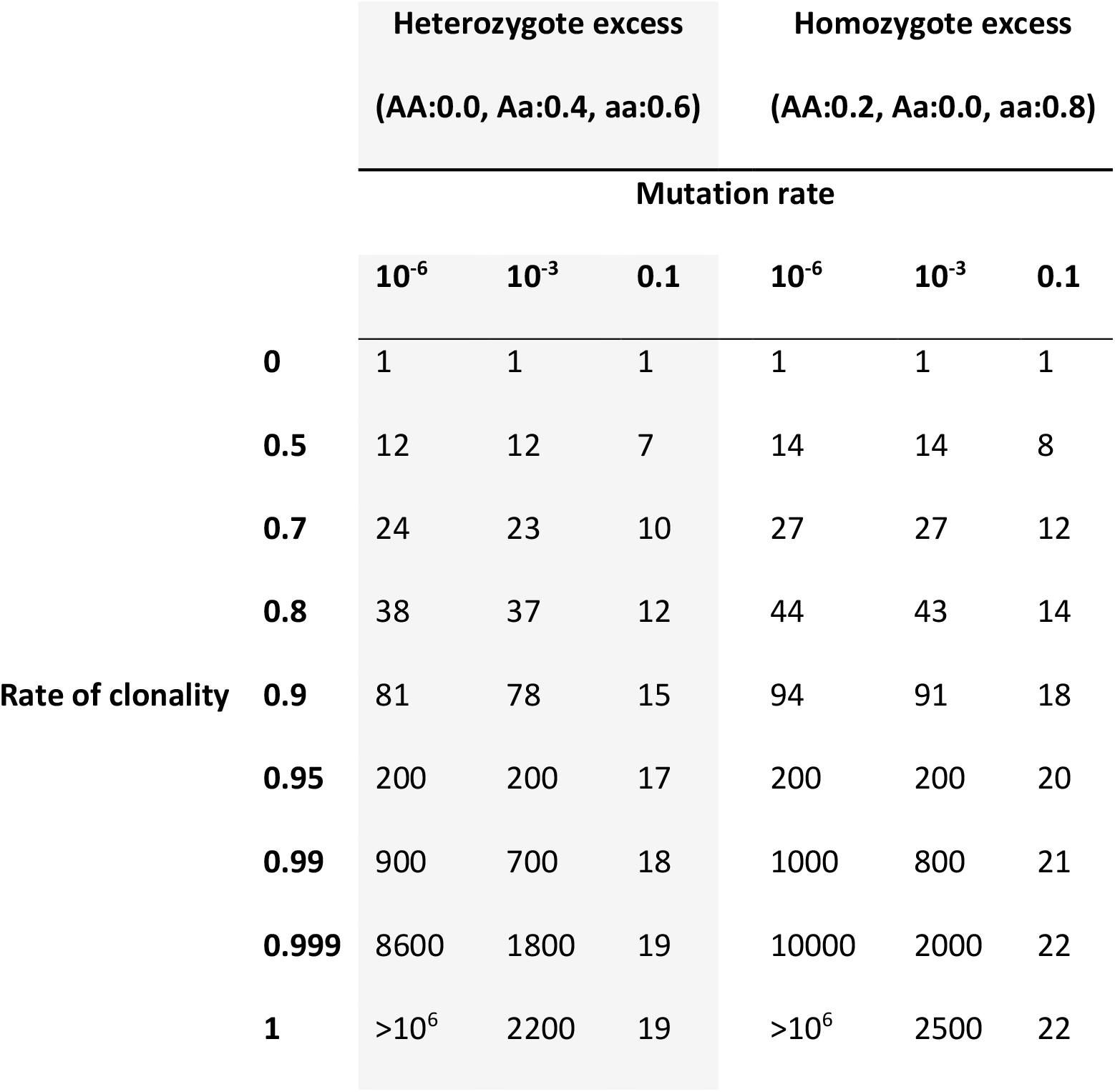
Number of generations required to reach Hardy-Weinberg proportions (*n*_*HW*_, eq.17) as function of clonality rate (row) and mutation rate (column), and two initial genotype frequency states, corresponding to initial heterozygote excess and homozygote excess (group of columns respectively in grey and white).

**Figure 8:**
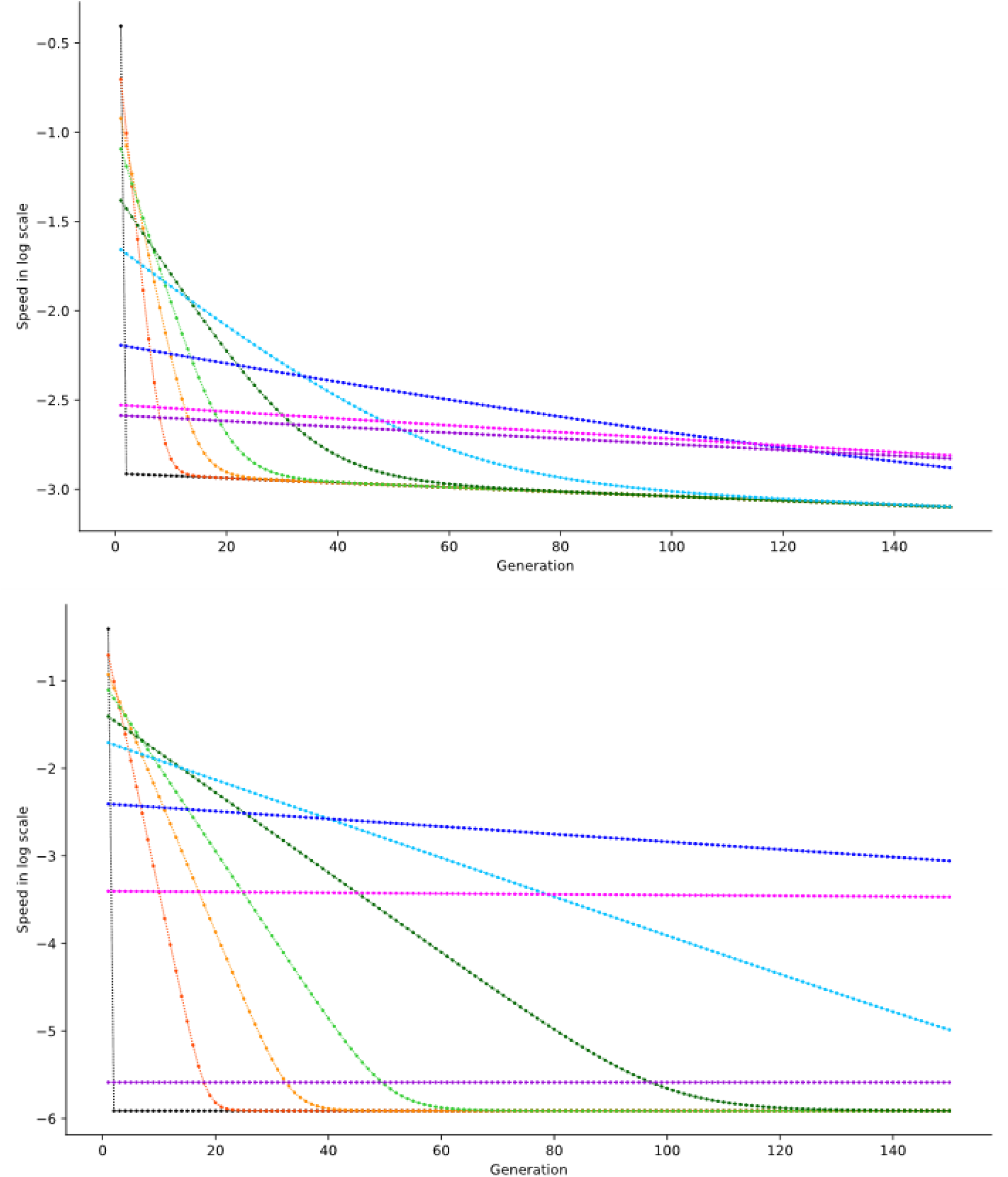
Speed along mean trajectories 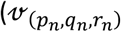, eq.14) originating from the initial point (aa:0.2, Aa:0, AA:0.8). A change in acceleration occurs upon reaching the Hardy-Weinberg parabola. The top panel corresponds to a mutation rate of 10^−3^ and the bottom panel to a mutation rate of 10^−6^. Line colors represent different clonality rates: black: c=0, orange: c=0.5, gold: c=0.7, limegreen: c=0.8, darkgreen: c=0.9, skyblue: c=0.95, blue: c=0.99, fuchsia: c=0.999, darkviolet: c=1.

Upon reaching the Hardy-Weinberg parabola, all populations, regardless of their clonality rate, adopt a common evolutionary pace dictated solely by mutation (Table 2). Again, increasing mutation accelerates the return to the stable equilibrium.

**Table 2.**
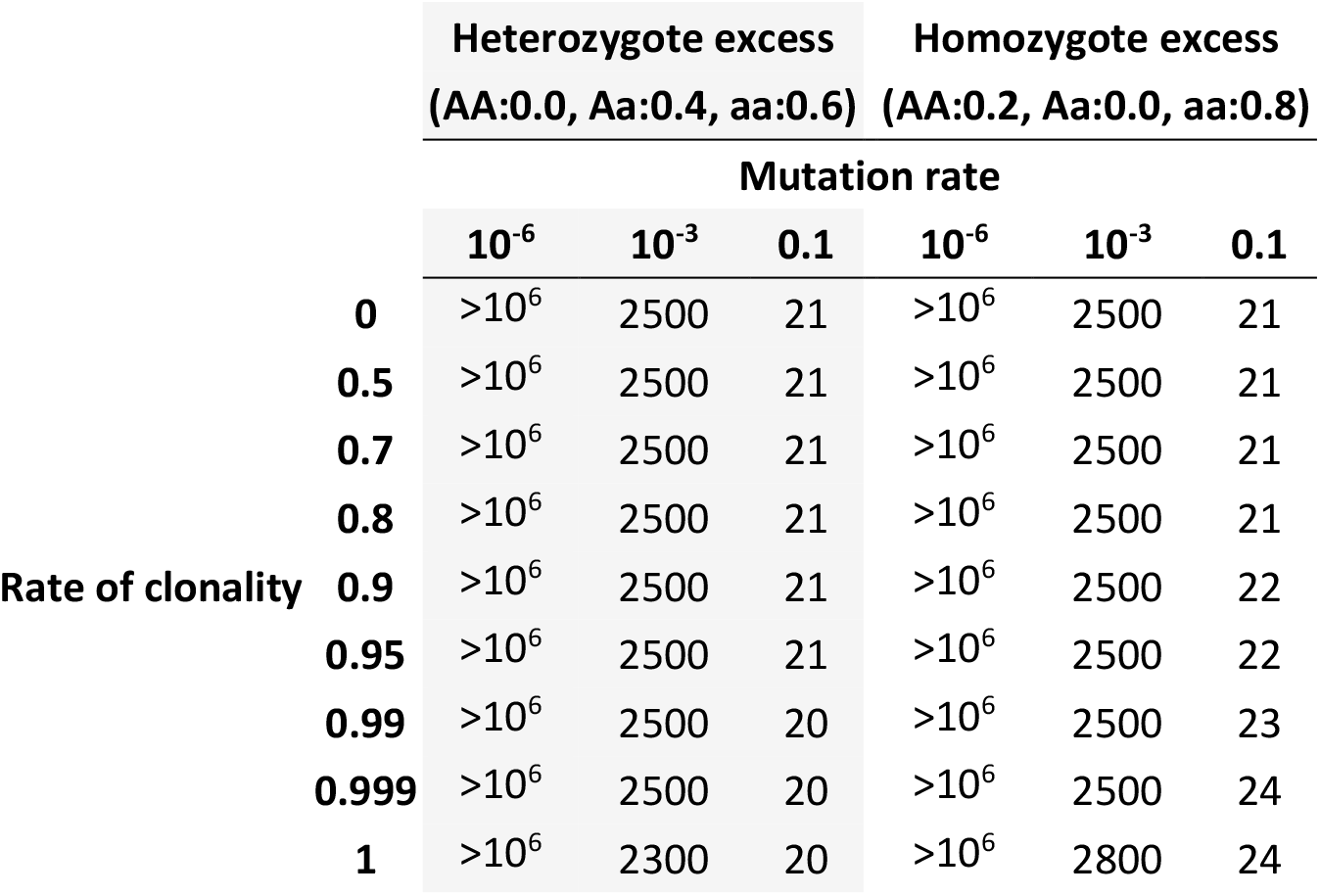
Number of generations required to reach equilibrium (*n*_*e*_, eq.18) as function of clonality rate (row) and mutation rate (column), and two initial genotype frequency states, corresponding to initial heterozygote excess and homozygote excess (group of columns respectively in grey and white).

Interestingly, from the same initial point, the higher the clonal rate, the further apart the trajectories iterate along the Hardy-Weinberg parabola (Figure 9). However, these deviations occur at such a small scale that they would require exceptionally precise genotype frequency estimates from empirical datasets to distinguish different rates of clonality. As trajectories approach the Hardy-Weinberg parabola and become aligned with it, a transient deceleration is observed before reaching the parabola, followed by a rebound once the system iterates along it (Figure S1).

**Figure 9:**
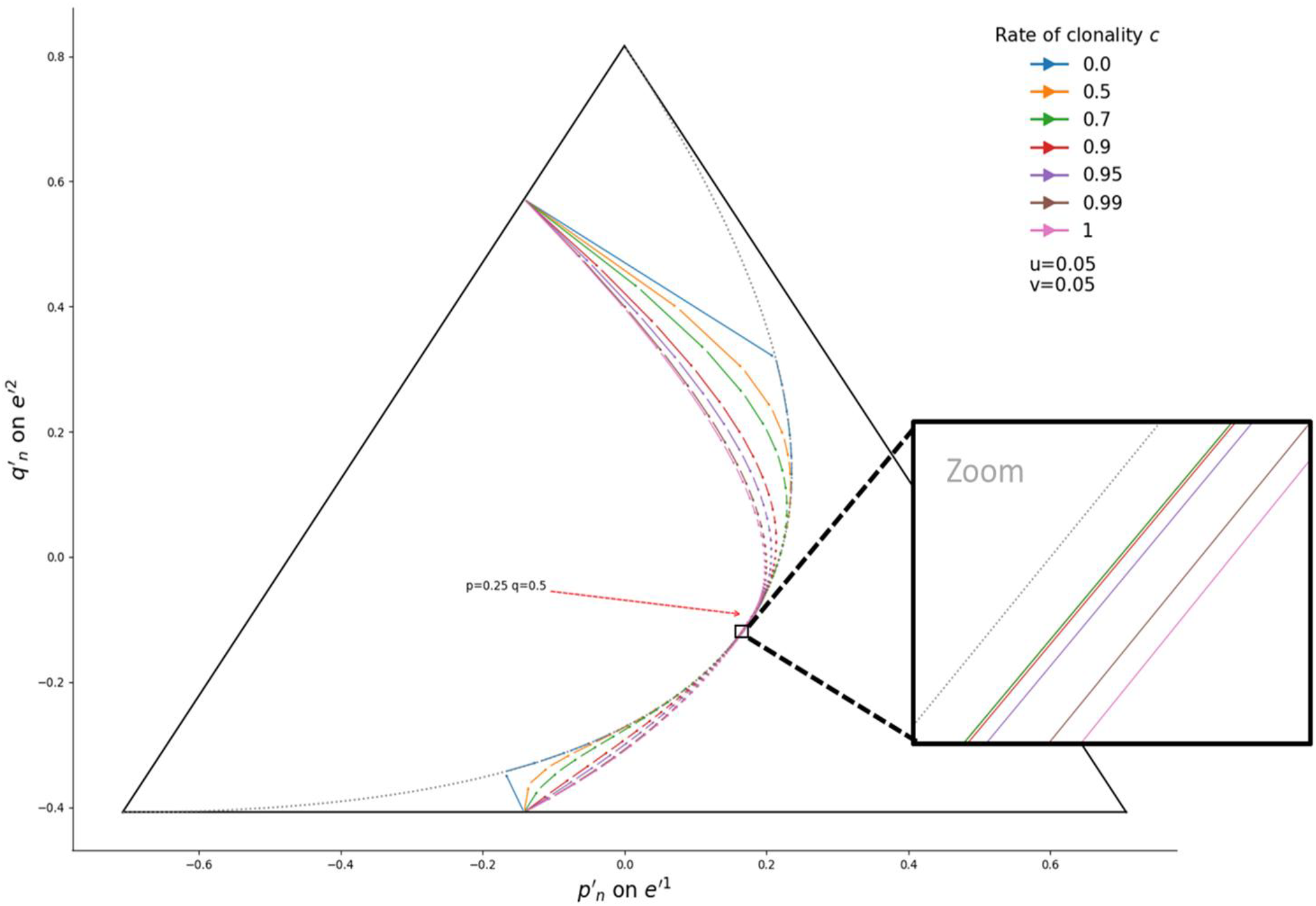
Trajectories of genotype frequencies along generations in a population reproducing with *c* = 0.9 and mutating with *u* = 0.05. Each arrow represents the change in genotype frequencies over one generation. Line colors indicate clonality rates (see legend, top right). The Hardy-Weinberg parabola is shown as a dashed grey line. All trajectories converge towards (AA:1/4, Aa:1/2, aa:1/4). Trajectories at bottom right originate from the same initial point (AA:0.6, Aa:0.4, aa: 0) while trajectories at top left originate from the same initial point (AA:0.2, Aa:0, aa: 0.8). The inset shows a close-up view near the stable equilibrium, showing that the higher the rates of clonality, the further trajectories iterate along Hardy-Weinberg parabola. In the inset, trajectories for *c* = 0, *c* = 0.5 and *c* = 0.7 appear to overlap, but would be distinct and ordered upon further magnification.

Because the number of generations required to reach the Hardy-Weinberg parabola increases with clonality, populations sharing identical initial conditions but reproducing with differing rates of clonality follow distinct trajectories: higher rates of clonality extend the first phase (return to Hardy-Weinberg parabola), enabling mutation to shift allele frequencies and redirect the sequence of distributions of genotype frequencies prior to convergence (Figure 9).

### 3.1 Concentration ellipse

Now considering the stochasticity of the trajectories in finite population sizes, concentration ellipses exhibit maximum size in the interior of the simplex, diminishing toward boundaries and vertices (Figure S2). Crucially, these ellipses are independent of clonality rate (see Appendix): both the area and axis lengths of the concentration ellipse depend solely on (*p*_*n*_, *q*_*n*_, *r*_*n*_) and population size *N*. Clonality rate in fact determines only which sequence of distributions of genotype frequencies the population traverses (Figure 9), by altering the mean speed and direction of the trajectory (Figures 7 and 8).

During convergence to Hardy-Weinberg proportions, ellipse area grows as the major axis extends disproportionately relative to the minor axis, peaking just before the parabola is reached, then decreasing slightly (Figures 4, S3 and S4). Once on the Hardy-Weinberg parabola, ellipse area resumes growth at a reduced rate, continuing until the stable equilibrium is attained. At and near the Hardy-Weinberg parabola vertex and the stable equilibrium, across all trajectories and regardless of clonality or mutation rates under reciprocal mutation between alleles, the concentration ellipses exhibit marked eccentricity, with the major axis oriented toward both heterozygote excess and homozygote excess (see Appendix).

### 3.2 Impacts on the dynamics of population genetic indices

Clonality rate does not alter allele frequencies along mean evolutionary trajectories (eq.6). As a result, gene diversity and its dynamics across generations do not differ among clonality rates (Figure S5). It is the relative balance between genetic drift and mutation rate (not clonality itself) that determines the mean gene diversity. In fact, clonality delays the return to Hardy-Weinberg expected heterozygosity levels (Figure S6), without altering the mean allele frequencies. This in turn alters the mean F_IS_ along generations (Figure S7).

In partially or fully clonal populations, transient departures toward positive or negative mean F_IS_ values are therefore expected only during the generations required for genotype frequencies to return to the Hardy-Weinberg parabola following a perturbation displacing them from Hardy-Weinberg proportions (Table 1). Once the Hardy-Weinberg parabola is reached, mean F_IS_ remains close to zero until the population reaches the stable equilibrium (Table 2). However, slight negative F_IS_ are still expected in finite population due to the concavity of the Hardy-Weinberg parabola (Figure S8) and, potentially, to the transient deviation of mean genotype frequencies from the parabola, depending on the nature of the preceding perturbation (zoom in, Figure S8).

## 4 Discussion

In this study, we developed a mathematical model that enables the computation of both the mean and the variance of genotype frequencies across successive generations within a given population with substantially greater speed and efficiency than proposed in Stoeckel and Masson (2014), even when transition matrices are approximated as sparse (Reichel et al. 2015). By reformulating the model from a discrete to a continuous representation of genotype frequencies, which is particularly appropriate for large population sizes, our approach facilitates the tracking of genotype and allele frequencies across generations. This reformulation also enables explicit characterization of the temporal evolution of the associated dispersions (concentration ellipses), thereby offering a fuller description of the stochastic variation in genotype frequency dynamics. We found that, for fixed population size and mutation rate, both different initial conditions and different clonality rates generate distinct, one-to-one invertible mean trajectories. These results suggest that it may be possible to infer both backward and forward mean genotype frequency trajectories, as well as the evolutionary forces driving them, from time-series genotyping data collected in monitored populations.

### 4.1 A common stable equilibrium and a global two-phase dynamic out of equilibrium

Our results further indicate that partially and fully clonal populations converge to the same stable equilibrium as fully sexual populations. This stable equilibrium only depends on the relative mutation rates between alleles. Out of equilibrium, genotype frequency trajectories follow two-phase dynamics: first, genotype frequencies return toward Hardy-Weinberg proportions and then evolve along the Hardy-Weinberg parabola until the equilibrium point is reached (corresponding to 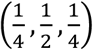 under reciprocal mutation rates *u* = *v*). In the first phase, convergence toward Hardy-Weinberg proportions decelerates as clonality rates increase. In the second phase, trajectories iterate precisely along the parabola in sexual populations but deviate increasingly from it as clonality rate increases. These two phases (return to Hardy-Weinberg proportions, and iteration along Hardy-Weinberg parabola towards the stable point) also occur in exclusively sexual populations; However, the first phase is greatly compressed, completing within a single generation. As clonality increases, populations take longer to return to Hardy-Weinberg proportions. During the time required to return to the Hardy-Weinberg parabola, mutation acts to alter allele frequencies, causing mean trajectories to pass through a different sequence of genotype frequency distributions (Figure 9).

### 4.2 Consequences for negative and positive F_IS_ in clonal populations

Our equations and results further indicate that variation in F_IS_ distributions with clonality arises because additional evolutionary forces (e.g., genetic drift, selection) and transient demographic or migration events drive genotype frequencies at certain loci away from equilibrium and beyond Hardy-Weinberg proportions. In simulation studies, departures tend toward negative F_IS_ because initial genetic diversity is often set to its maximum (Marshall and Weir 1979; Balloux et al. 2003, Stoeckel et al. 2021a,b) or due to non-equilibrium demographic scenarios (Berg and Lascoux 2000). When examining genotype frequency distributions at steady-state equilibrium, or after a large number of generations regardless of initial conditions, a prevalence of negative F_IS_ is indeed observed: Because the Hardy-Weinberg parabola is concave toward the region of homozygote excess, when populations are near their equilibrium and Hardy-Weinberg parabola, the area of the concentration ellipse above it (corresponding to slight heterozygote excess) is larger than the area below it (corresponding to slight homozygote excess; Figure S9), further aided by the eccentricity of the ellipse, whose major axis extends perpendicular to the parabola. As these ellipses do not depend on clonality (see Appendix), these preferential departures toward negative F_IS_ are also expected in fully sexual populations, particularly when population size is small. However, in that case, these deviations are very transient, as they typically persist for only a single generation, unless genetic drift or any other dominant evolutionary force impede the return to Hardy-Weinberg proportions. In partially clonal populations, by contrast, such deviations persist on average for multiple generations before genotype frequencies return to Hardy-Weinberg proportions. Moreover, these deviations may drift slightly further in small population sizes, as this extended period allows further deviations to accumulate.

In addition, as population size decreases, the area of the concentration ellipse increases, increasing the probability of observing loci departing from Hardy-Weinberg proportions, including towards negative F_IS_ since, in partially clonal populations, genetic drift operates at the genotype rather than the allele scale. Under clonal reproduction, genetic drift (more aptly termed *genotype drift* in this context) can therefore fix any genotype, including heterozygous, the number of which *n*_*Het*_ increases with the number of alleles as *n*_*Het*_ = (*k* − 1)!. Thus, *F*_*IS*_ = −1 is expected only when a single heterozygous genotype becomes fixed through genotype drift under a low mutation rate. The probability of this outcome is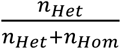where 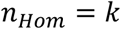, This probability reaches its maximum under the infinite-sites mutation model.

Altogether, these processes explain why the proportion of slightly negative and positive F_IS_ values constitutes a reliable indicator of clonality when genetic drift dominates, but becomes less reliable when mutation rates are comparable to or exceed the strength of genetic drift (Stoeckel and Masson 2014; Reichel et al. 2016; Rouger et al. 2016) and why, ultimately, variance of F_IS_ values among genotyped loci remains one of the most robust signals for inferring clonality rates from a single-sample population survey (Arnaud-Haond et al. 2020; Stoeckel et al. 2021a,b).

Our results even suggest that, for known population size and mutation rates, the absolute deviations of observed genotype frequencies from Hardy-Weinberg proportions, rather than negative F_IS_ (heterozygote excess) alone, may provide a better proxy for estimating clonality rates in polyploid populations. Nonetheless, the balance between positive and negative deviations remains useful for distinguishing clonality from selfing, as selfing is expected to generate predominant excesses of homozygosity across loci. In population monitoring, the number of successive generations required for genotype frequencies, displaced from Hardy-Weinberg proportions at each marker, to return to these proportions offers a promising avenue for inferring clonality rates.

### 4.3 Perspectives

Finally, we also established that mean evolutionary trajectories are invertible and bijective, enabling inference of clonality rates across multiple generations without requiring samples to be genotyped at strictly one-generation intervals as in current methods (Becheler et al. 2017 for diploids; Stoeckel et al. 2024 generalization for any autopolyploids). Using reciprocal mutation rates between alleles, i.e., ∀ (*i, j*), *u*_*ij*_ = *u*, produce symmetric dynamics about the axis of symmetry passing through the Hardy-Weinberg vertex, 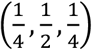 for a diallelic locus, bisecting the parabola into mirror-image halves. Under non-reciprocal mutation rates (*u*_*ij*_ ≠ *u*_*ji*_), this symmetry is broken and the stable equilibrium shifts away from the Hardy-Weinberg parabola vertex. Nevertheless, the fundamental structure of the system remains: genotype frequency trajectories converge toward a fixed point that we conjecture to always lie on the Hardy-Weinberg parabola, regardless of clonality rate. Incorporating non-reciprocal mutation rates between alleles may therefore be necessary to apply our model to empirical datasets, particularly when SNP mutations exhibit asymmetric transition rates (e.g., G/C↔A/T mutation biases). Our generalization to multiple alleles with non-reciprocal mutation rates and the possibility to include population size changing over time (see 2.1) lay the groundwork for developing inference methods applicable to empirical experiments and long-term population monitoring.

## Statements and Declarations

The authors declare no competing interests.

## CRediT authorship contribution statement

### Solenn Stoeckel

Conceptualization, Formal analysis, Funding acquisition, Investigation, Methodology, Project administration, Computation, Software, Validation, Visualization, Writing – original draft, Writing – review and editing. **Jean-Pierre Masson**: Conceptualization, Formal analysis, Investigation, Methodology, Computation, Validation, Visualization, Writing – original draft, Writing – review and editing.

## Appendix

Technical aspects considering concentration ellipses

## Supplementary information

Supplementary illustrative figures that support results, discussion and conclusions.

## Data availability

Code of mathematical developments and reproducible results can be assessed at https://forge.inrae.fr/solenn.stoeckel/dygenclon

## Appendix

### Technical aspects considering concentration ellipses

For one population size *N* at generation *n*

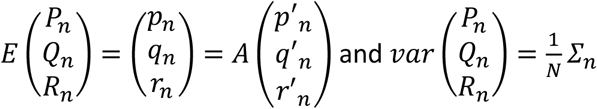

with 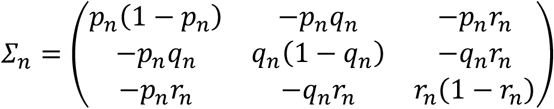

Remember that

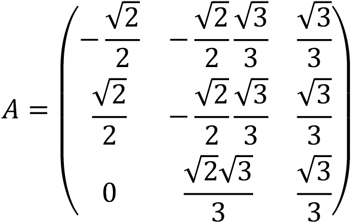

A principal component analysis gives two non-zero eigen values 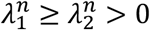 and two associated eigenvectors 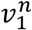 and 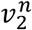

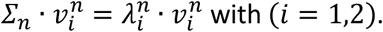

To simplify the notation going forward, we now omit the superscript index *n*, which indicates the generation.

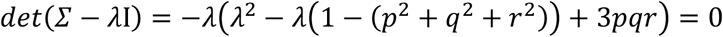

*det*(*Σ* − *λ*I) = −*λ*(*λ*^2^ − *λ*(1 − (*p*^2^ + *q*^2^ + *r*^2^)) + 3*pqr*) = 0

The non-zero eigenvalues are solutions to *λ*^2^ − *λ*(1 − (*p*^2^ + *q*^2^ + *r*^2^)) + 3*pqr* = 0. On this characteristic polynomial, we identify

- the sums of the roots: *λ*_1_ + *λ*_2_ = 1 − (*p*^2^ + *q*^2^ + *r*^2^);
- the product of the roots: *λ*_1_*λ*_2_ = 3*pqr*;

Hence the explicit formula for eigenvalues:

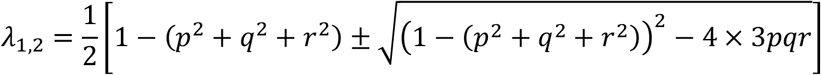

or similarly

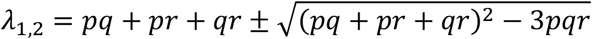

The eigenvectors *v*_1_ and *v*_2_ are orthogonal and indicate directions of the two axes of the concentration ellipse in the plane Π^*^.

From *Σ* and *λ*_1_, we obtain the eigenvector 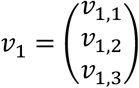 associated with *λ*_1_:

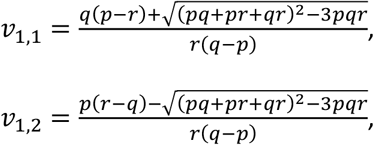

and *v*_1,3_ = 1.

From *Σ* and *λ*_2_, we obtain the eigenvector 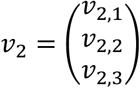 associated with *λ*_2_:

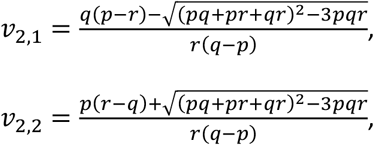

and *v*_2,3_ = 1.

The eigenvector *v*_1_ (respectively *v*_2_) gives the direction of the major (respectively minor) axis of the concentration ellipse.

For the graphical representation, we express them in the new basis *e*_1_′, *e*_2_′, *e*_3_′:

*v*′_*i*_ = *A*^*t*^ · *v*_*i*_ with *i* = 1,2 and 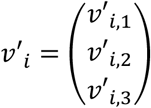

More concretely:

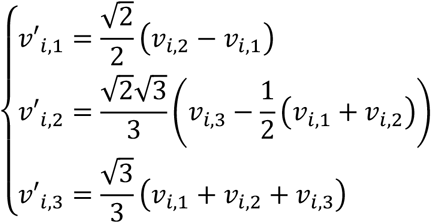

*v*′_1_ and *v*′_2_ are orthogonal vectors and they belong to Π^*^; So *v*′_*i*,3_ = 0. Therefore, for a population size *N*, the reference ellipse is defined as follows:

In the basis *e*_1_′, *e*_2_′, *e*_3_′, *v*′_1_ (respectively *v*′_2_) gives the direction of the major (respectively minor) axis. 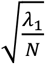 (respectively 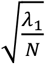 ) gives the length of the semi-major (respectively semi-minor) axis. The semi-major and semi-minor axes are equal for (*p* = ⅓, *q* = ⅓, *r* = ⅓), a point out of the Hardy-Weinberg proportions.

In the subspace Π^*^ generated by the vectors *e*_1_′ and *e*_2_′, the eigenvector *v*_1_ becomes *v*′_1_: 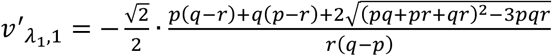,

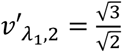and 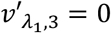

And the angle 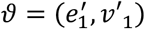 of the major axis with 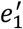 is given by:

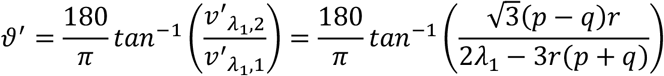

Remark: with the same approach, 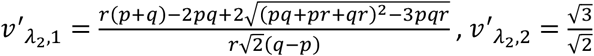, and 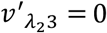 .

In the graphical representations, we choose a suitable scale for displaying the ellipses, disregarding population size. For a population of size *N*, a scaling by a factor of 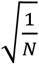 should be applied. This concentration ellipse associated with (*p*_*n*_, *q*_*n*_, *r*_*n*_) reduces to the only (*p*_*n*_, *q*_*n*_, *r*_*n*_) point for an infinite population size (*N* = ∞).

**Figure.**
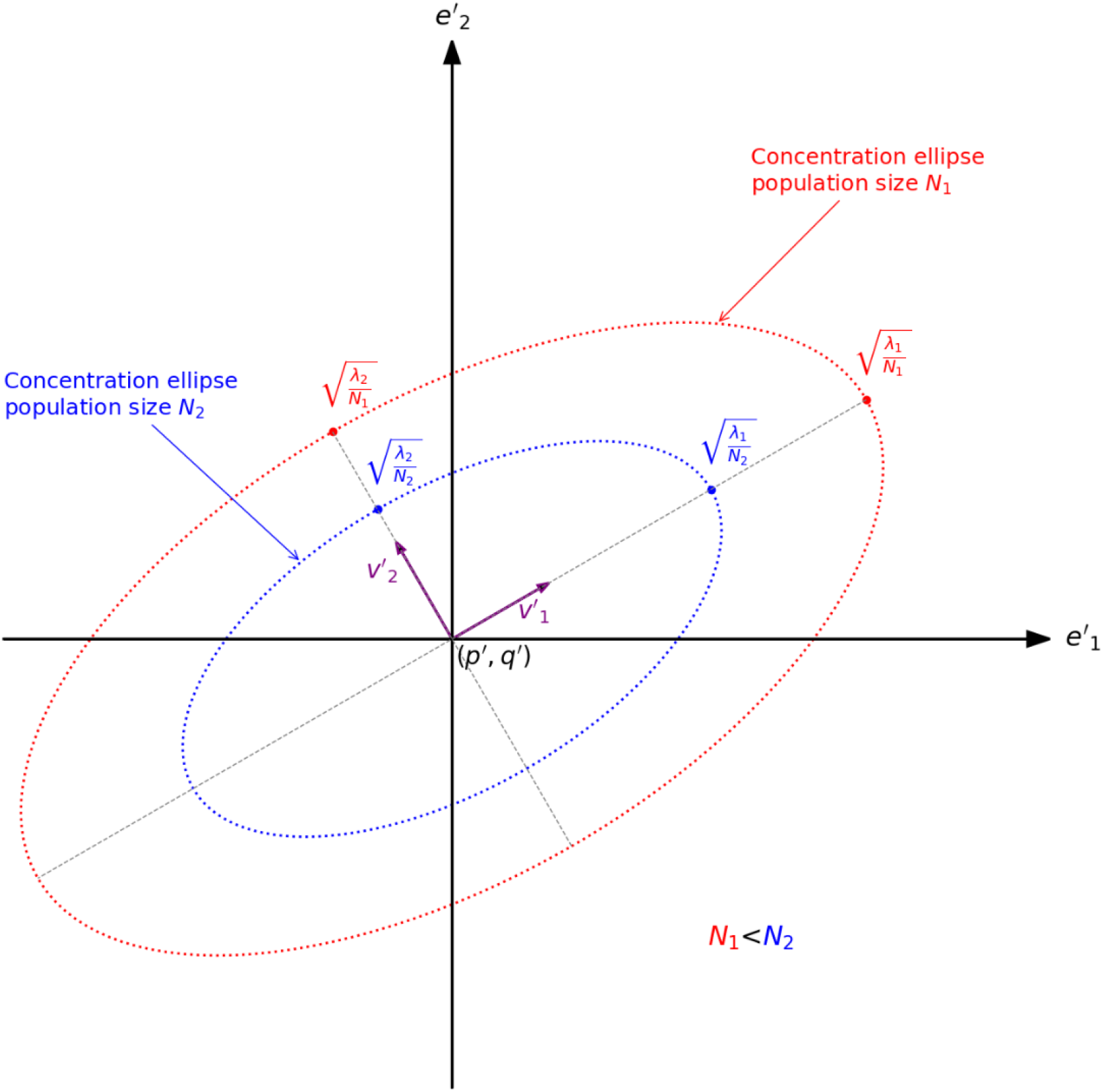
Concentration ellipses from a genotype frequency triplet (*p*_*n*_, *q*_*n*_, *r*_*n*_) for a same *k*^2^and scaling for two population sizes (*N*_1_ < *N*_2_) in the new basis *e*_1_′, *e*_2_′. In red, the concentration ellipse for a population size *N*_1_, and in blue for a population size *N*_2_. Are reported the semi-major lengths 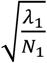 for *N* and 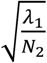 for *N*_2_ in the first axis given *v*′_1_ (grey dashed line on the purple arrow *v*′_1_ ) and semi-minor length 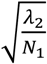 and 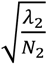 for *N*_2_ in the second axis given *v*′_2_ (grey dashed line on the purple arrow *v*′_2_).

### Accessing confidence regions around genotype frequencies

It is tempting to build a rectangular box using marginal confidence intervals. (*P, Q, R*) are approximately normally distributed with mean (*p, q, r*) and variances 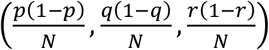 . It is reasonable to choose a risk 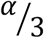 (Bonferroni method) to ensure an overall risk of *α*.

For example, for *α* = 5%, 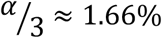 then:

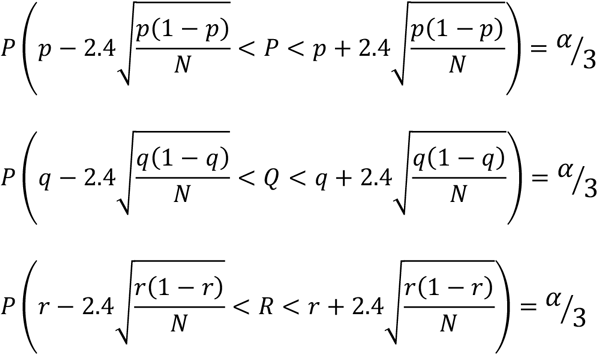

However, we must not forget that *p* + *q* + *r* = 1 and *P* + *Q* + *R* = 1

In the spirit of our developments, it is relevant to build confidence regions from concentration ellipses. *v*′_1_ and *v*′_2_ are now assumed to be normalized.

In the coordinate system *v*′_1_, *v*′_2_ the pair of independent random variable *X*^′^ − *p*′, *Y*^′^ − *q*′ has density *φ*(*x, y*) and 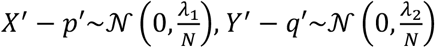

It is well known that:

- 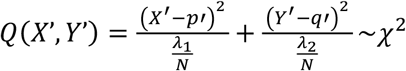with two degrees of freedom.
- 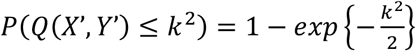

So, the confidence regions are then of the form

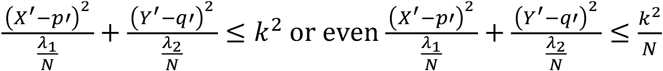

For example, for *α* = 5%, *P*(*Q*(*P*’, *Q*’) ≤ 6) ≃ 95%

We now represent these regions in the original space

a. Let 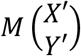 in (*e*_1_′, *e*_2_′) that is centered at 0. Let 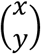 in (*v*′_1_, *v*′_2_) that is centered at 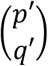 in (*e*_1_′, *e*_2_′). Then 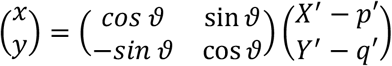 *v*′_1_ and *v*′_2_ are now assumed to be normalized. So *v*′_1,1_ = cos θ and *v*′_1,2_ = *sin* θ
b. 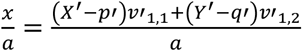 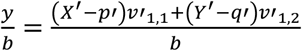 with 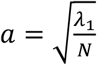 and *b* = 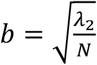 . The confidence region is now written:

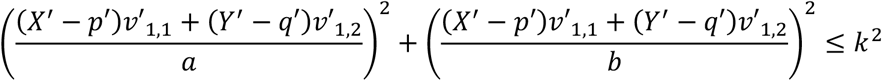
c. 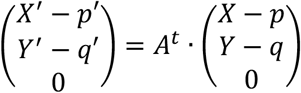 As *A*^*t*^ is orthogonal, we obtain the following relationships between the scale products:

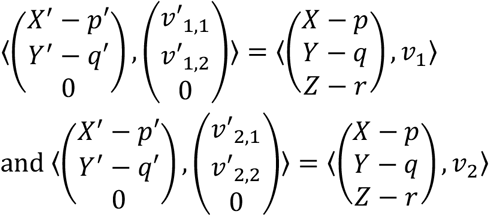

where *v*′_1_, *v*′_2_, *v*_1_, *v*_2_ are supposed to be normed in these writings.

So (*X*^′^ − *p*′)*v*′_*i*,1_ + (*Y*^′^ − *q*′)*v*′_*i*,2_ = (*X* − *p*)*v*_*i*,1_ + (*Y* − *q*)*v*_*i*,2_ + (*Z* − *r*)*v*_*i*,3_ with *i* = 1,2

In fine, a confidence region is the interior of the intersections of an ellipsoid and the plane *p* +

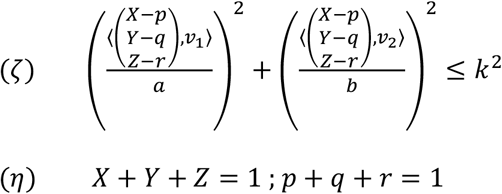

It is well known that the intersection of ellipsoid (*ζ*) and plan (*η*) is an ellipse as it was already seen before.

## Supplementary information

**Figure S1:**
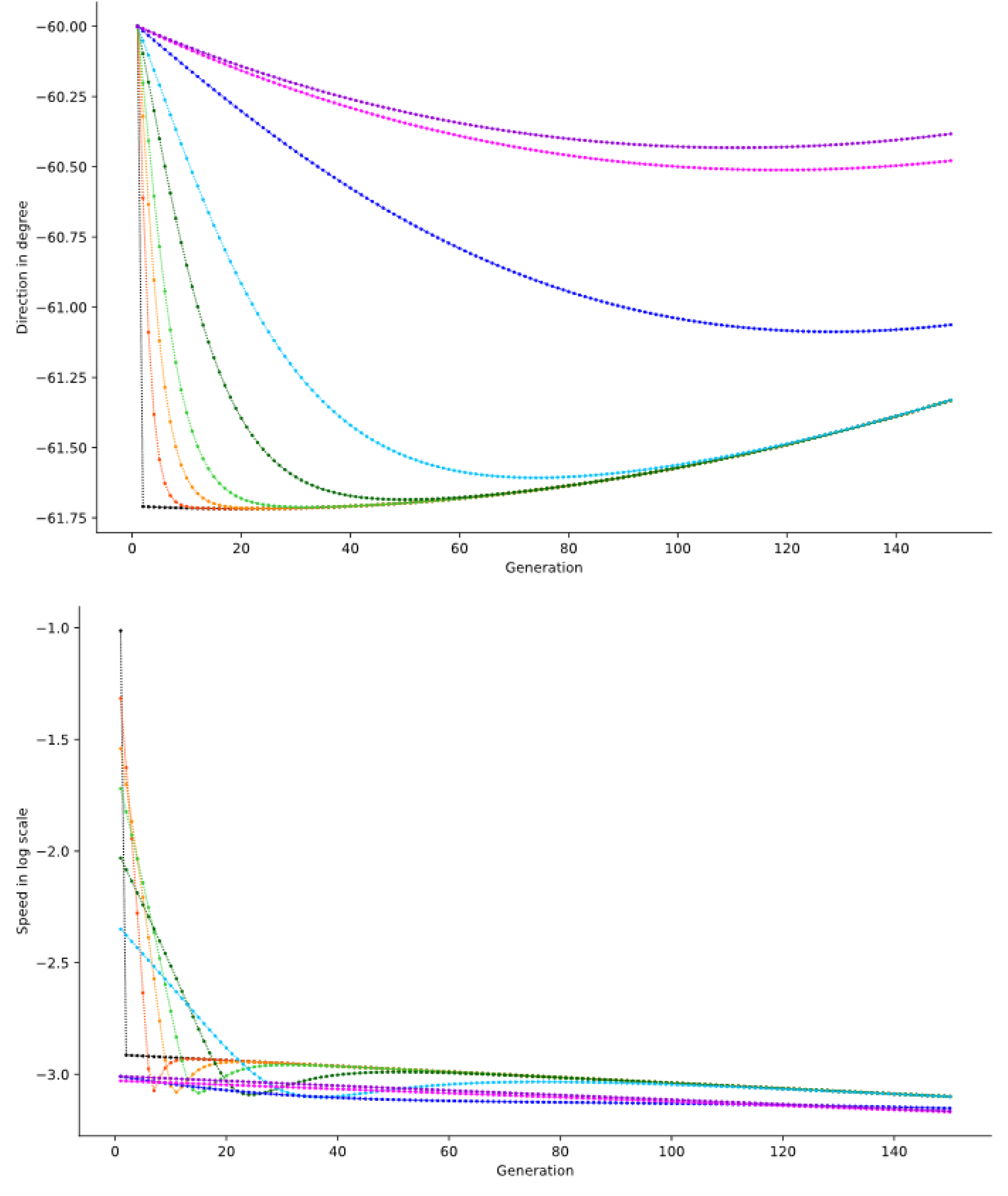
Angular direction and speed of trajectories originating from the initial point (aa:0, Aa:0.4, AA:0.6) with a mutation rate of 10^−3^. When a trajectory approaches the Hardy-Weinberg parabola tangentially owing to its initial direction, a velocity rebound is observed as trajectories begin to iterate along the Hardy-Weinberg parabola immediately after reaching it. Line colors represent different clonality rates: black: c=0, orange: c=0.5, gold: c=0.7, limegreen: c=0.8, darkgreen: c=0.9, skyblue: c=0.95, blue: c=0.99, fuchsia: c=0.999, darkviolet: c=1.

**Figure S2:**
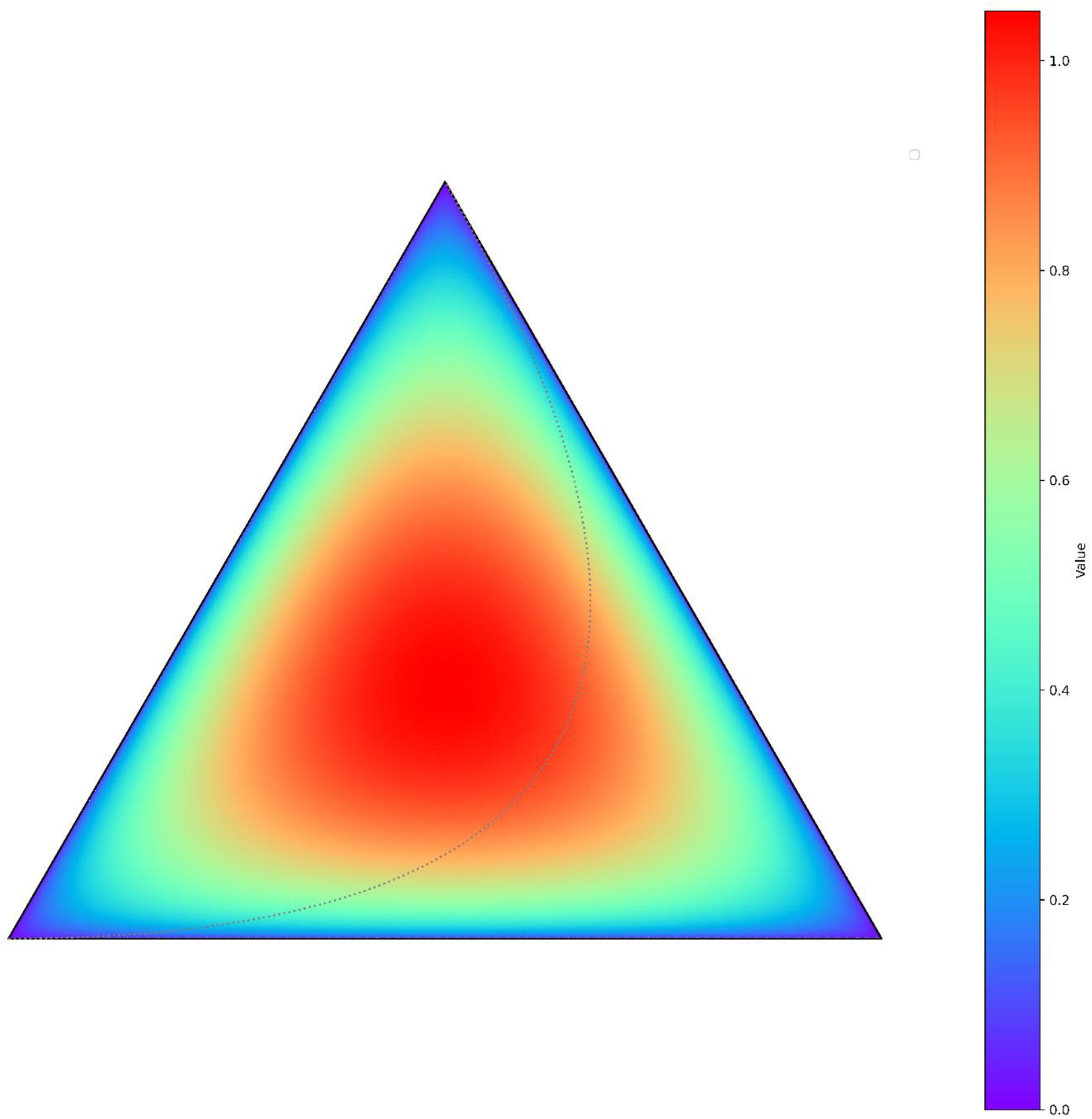
Heatmap of the concentration ellipse areas *A* (eq.15) across the subspace Π^*^, for a fixed population size *N* = 1000 and a reciprocal mutation rate *u* = 0.001 between the two alleles. The correspondence between colors and area values is indicated in the color bar. In grey dotted line, the Hardy-Weinberg parabola in the same subspace.

**Figure S3:**
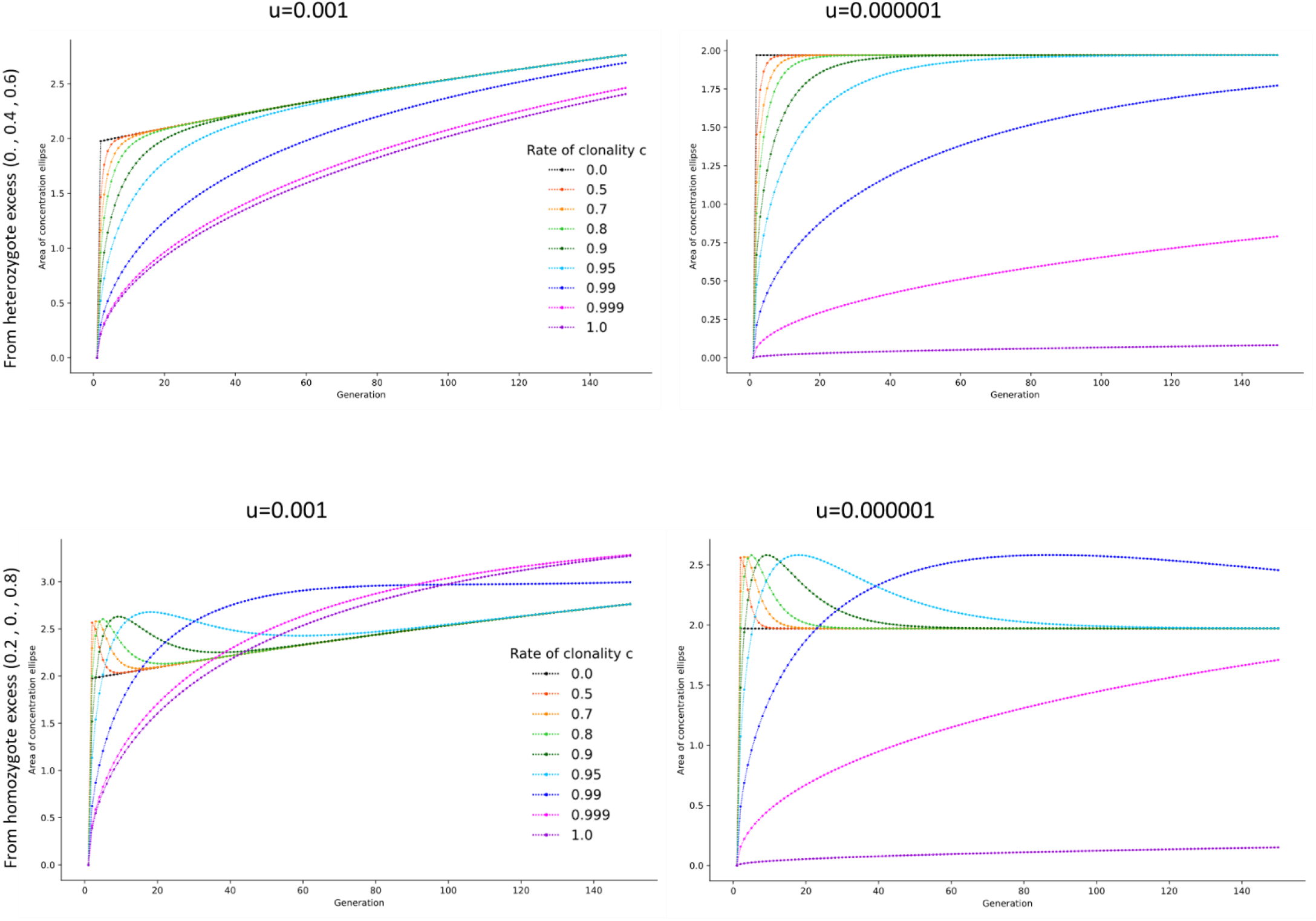
Dynamics of the concentration ellipses areas (eq. 15) across generations. First row: initial genotype frequencies corresponding to heterozygote excess (aa:0, Aa:0.4, AA:0.6); Second row: initial genotype frequencies corresponding to homozygote excess (aa:0.2, Aa:0, AA:0.8). Columns correspond to two reciprocal mutation rates: 10^−3^ (left) and 10^−6^ (right). Line colors represent different clonality rates: Black: c=0, orange: c=0.5, gold: c=0.7, limegreen: c=0.8, darkgreen: c=0.9, skyblue: c=0.95, blue: c=0.99, fuchsia: c=0.999, darkviolet: c=1.

**Figure S4:**
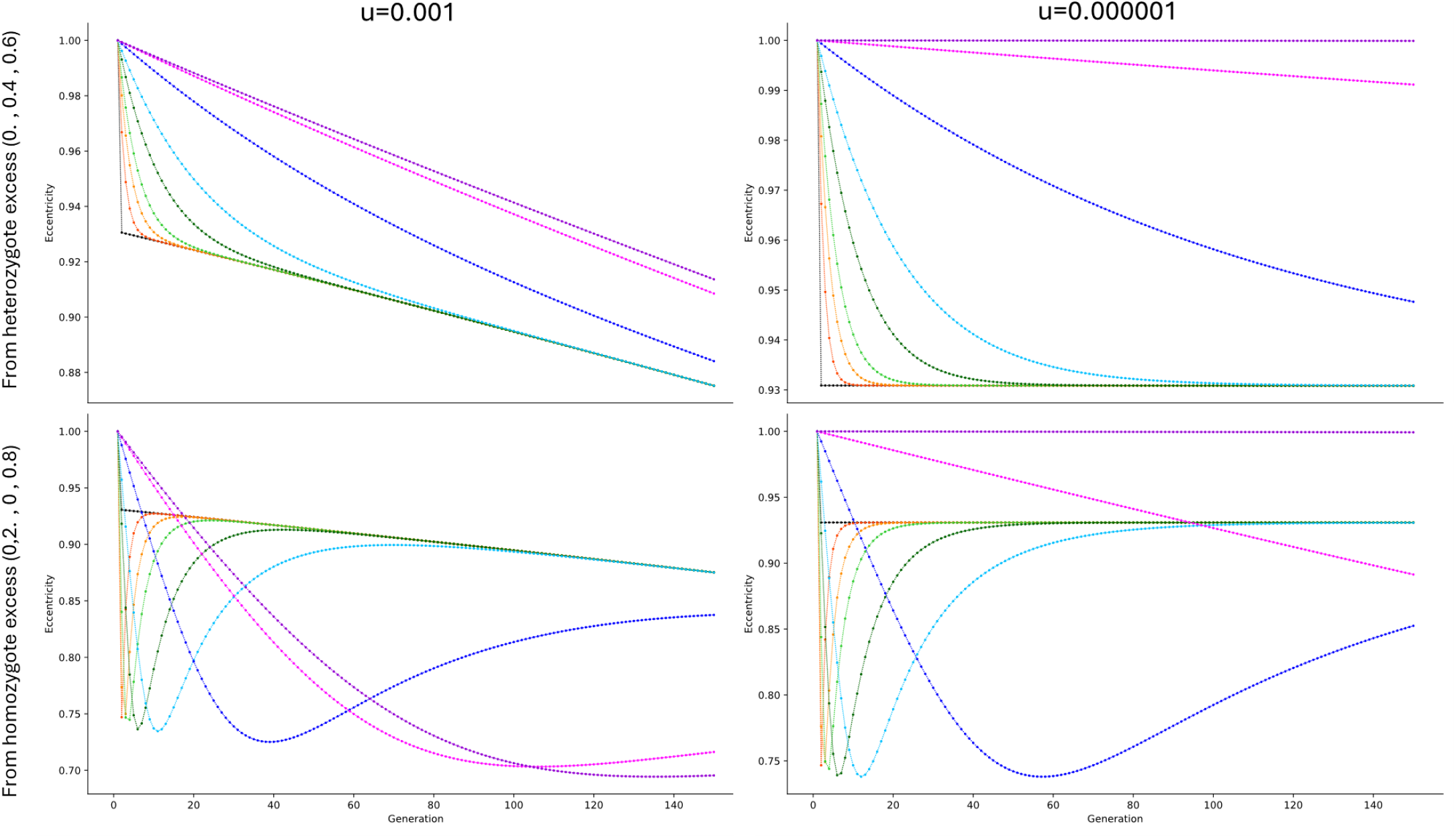
Dynamics of the concentration ellipses eccentricities (eq. 16) across generations. First row: initial genotype frequencies corresponding to heterozygote excess (aa:0, Aa:0.4, AA:0.6); Second row: initial genotype frequencies corresponding to homozygote excess (aa:0.2, Aa:0, AA:0.8).Columns correspond to two reciprocal mutation rates: 10^−3^ (left) and 10^−6^ (right). Line colors represent different clonality rates: Black: c=0, orange: c=0.5, gold: c=0.7, limegreen: c=0.8, darkgreen: c=0.9, skyblue: c=0.95, blue: c=0.99, fuchsia: c=0.999, darkviolet: c=1.

**Figure S5:**
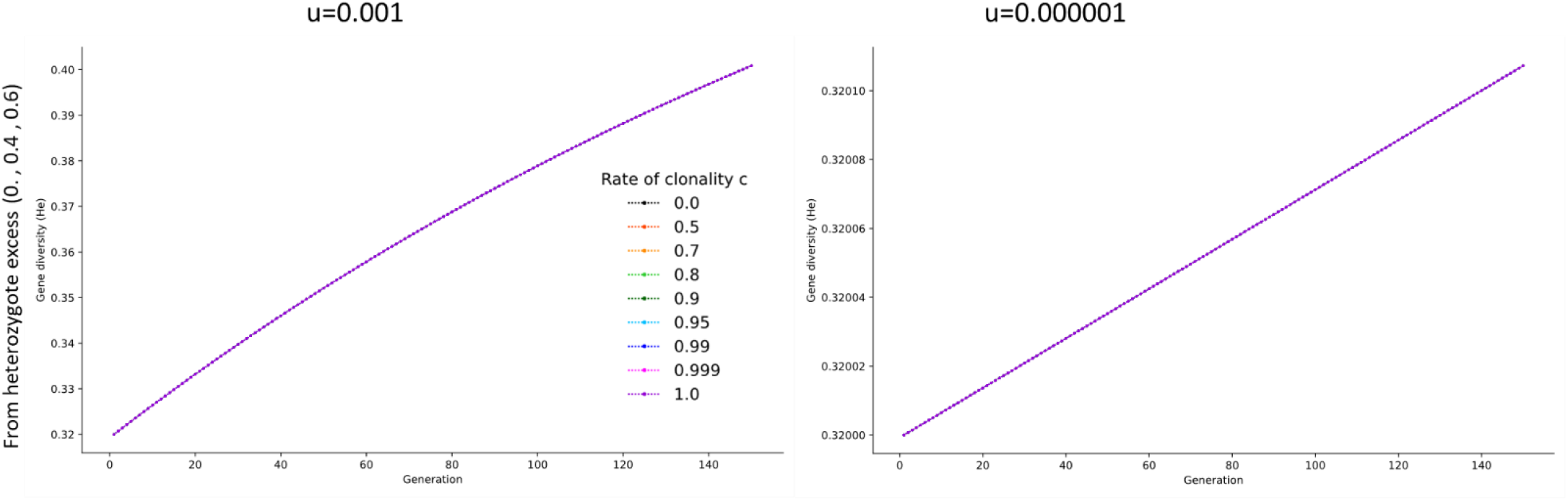
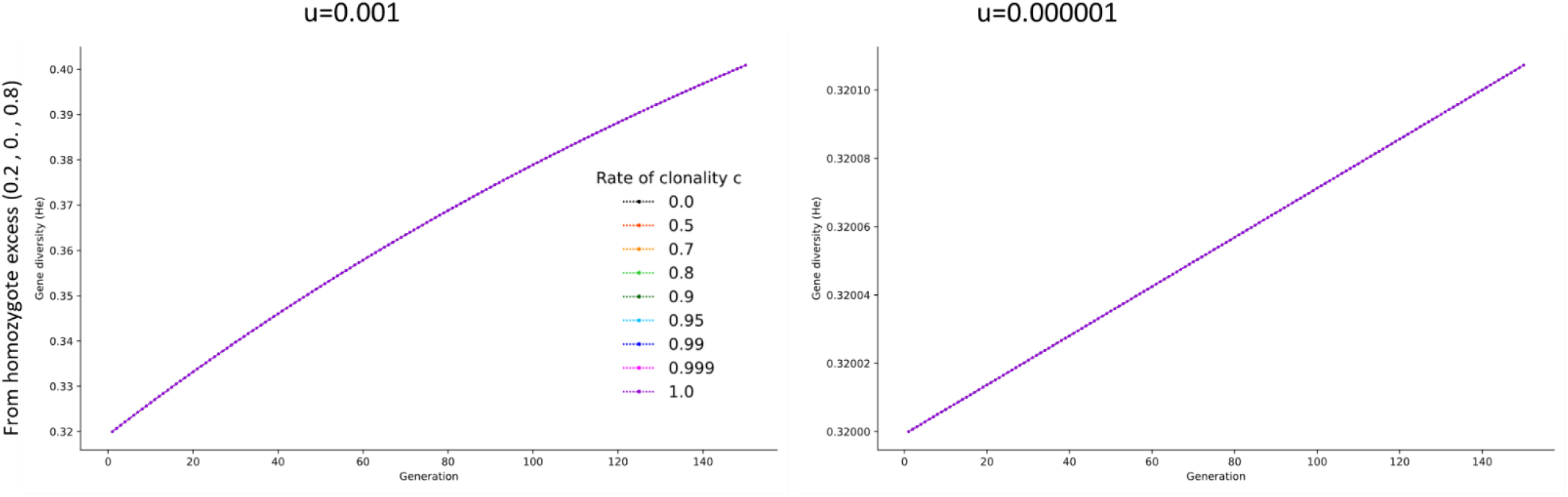
Dynamics of the gene diversity (He) across generations. First row: initial genotype frequencies corresponding to heterozygote excess (aa:0, Aa:0.4, AA:0.6); Second row: initial genotype frequencies corresponding to homozygote excess (aa:0.2, Aa:0, AA:0.8). Columns correspond to two reciprocal mutation rates: 10^−3^ (left) and 10^−6^ (right). Line colors represent different clonality rates: Black: c=0, orange: c=0.5, gold: c=0.7, limegreen: c=0.8, darkgreen: c=0.9, skyblue: c=0.95, blue: c=0.99, fuchsia: c=0.999, darkviolet: c=1.

**Figure S6:**
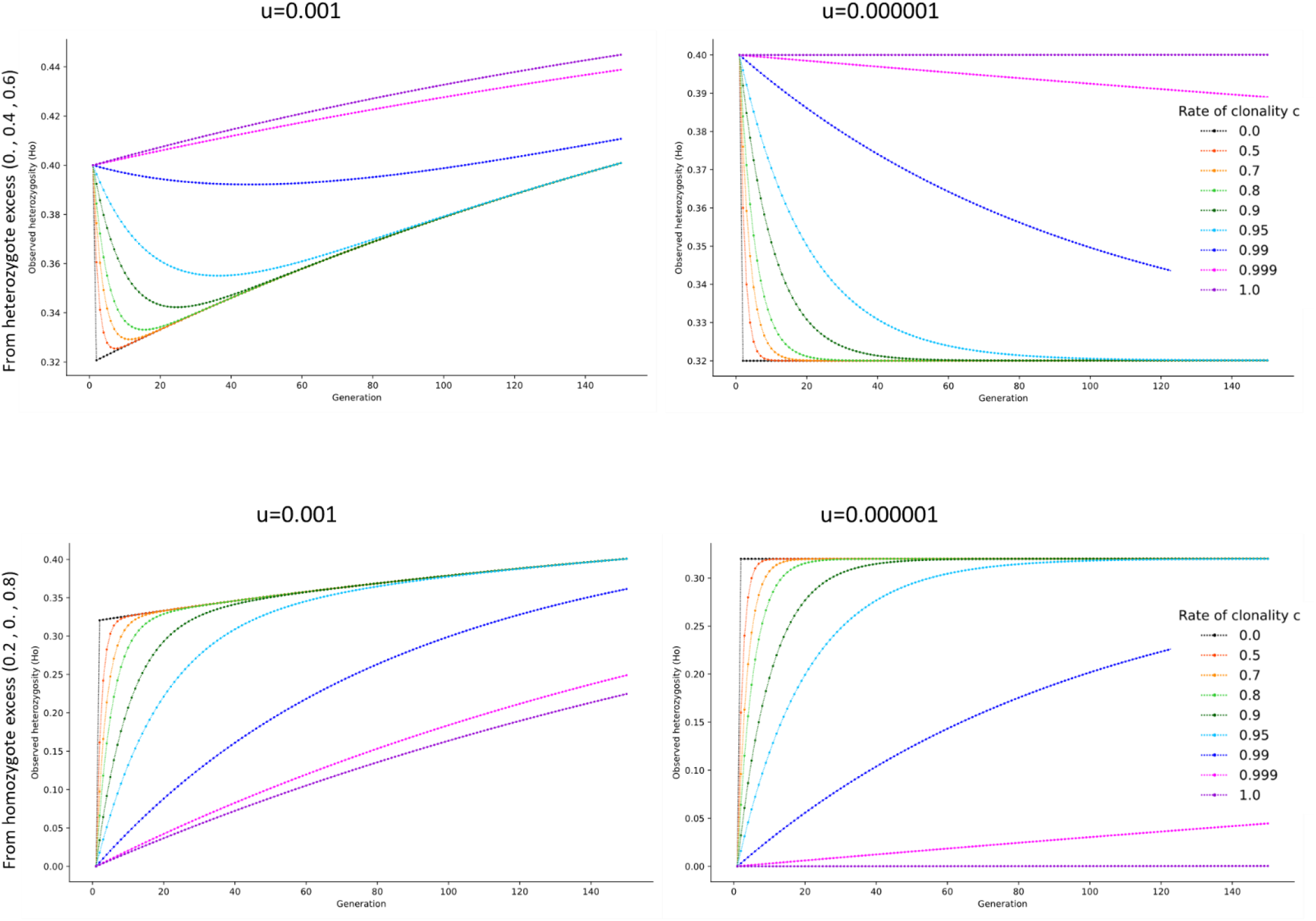
Dynamics of the observed heterozygosity (Ho, also *q* in equations) across generations. First row: initial genotype frequencies corresponding to heterozygote excess (aa:0, Aa:0.4, AA:0.6); Second row: initial genotype frequencies corresponding to homozygote excess (aa:0.2, Aa:0, AA:0.8). Columns correspond to two reciprocal mutation rates: 10^−3^ (left) and 10^−6^ (right). Line colors represent different clonality rates: Black: c=0, orange: c=0.5, gold: c=0.7, limegreen: c=0.8, darkgreen: c=0.9, skyblue: c=0.95, blue: c=0.99, fuchsia: c=0.999, darkviolet: c=1.

**Figure S7:**
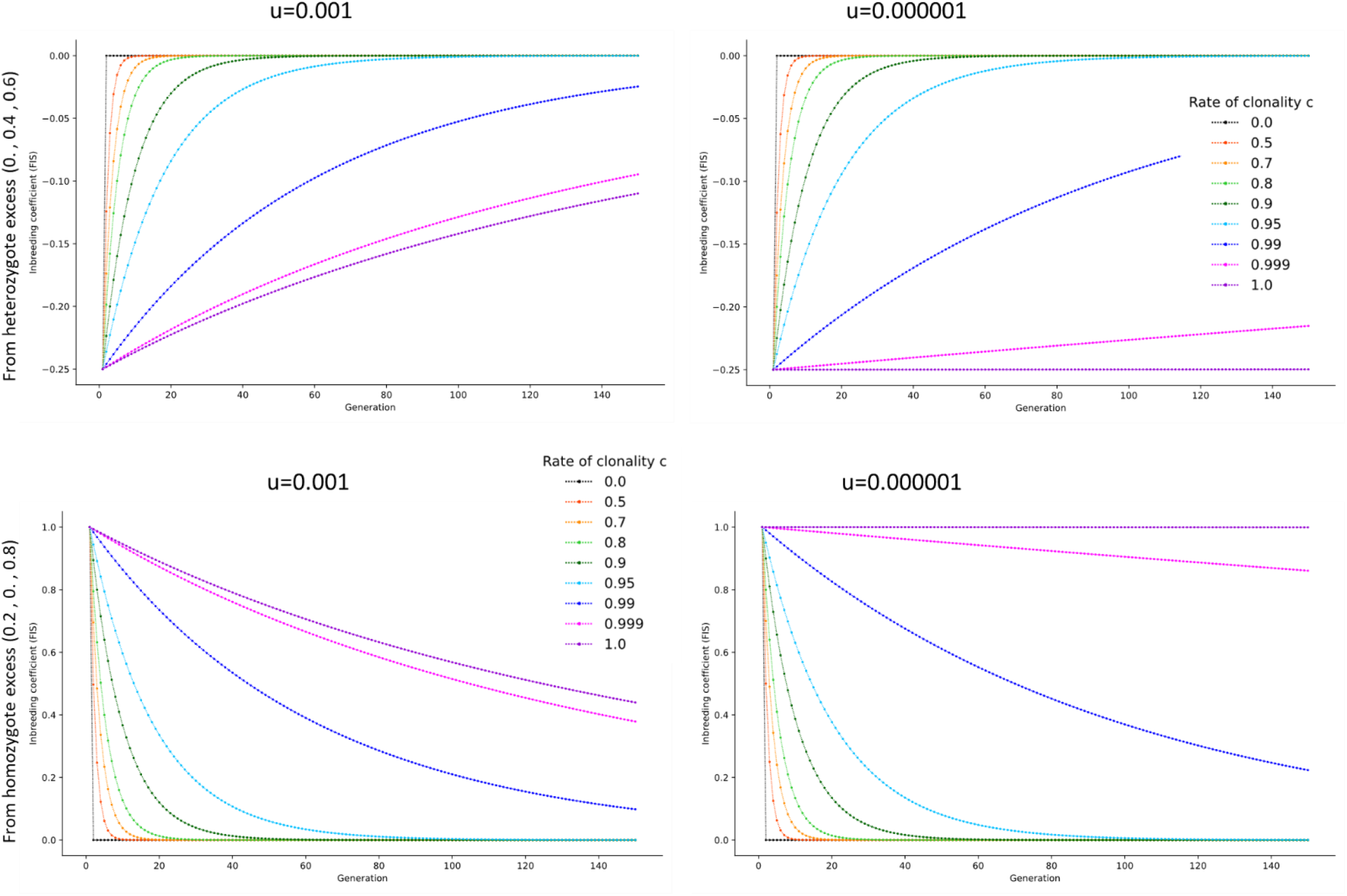
Dynamics of the inbreeding coefficient (F_IS_) across generations. First row: initial genotype frequencies corresponding to heterozygote excess (aa:0, Aa:0.4, AA:0.6); Second row: initial genotype frequencies corresponding to homozygote excess (aa:0.2, Aa:0, AA:0.8). Columns correspond to two reciprocal mutation rates: 10^−3^ (left) and 10^−6^ (right). Line colors represent different clonality rates: Black: c=0, orange: c=0.5, gold: c=0.7, limegreen: c=0.8, darkgreen: c=0.9, skyblue: c=0.95, blue: c=0.99, fuchsia: c=0.999, darkviolet: c=1.

**Figure S8:**
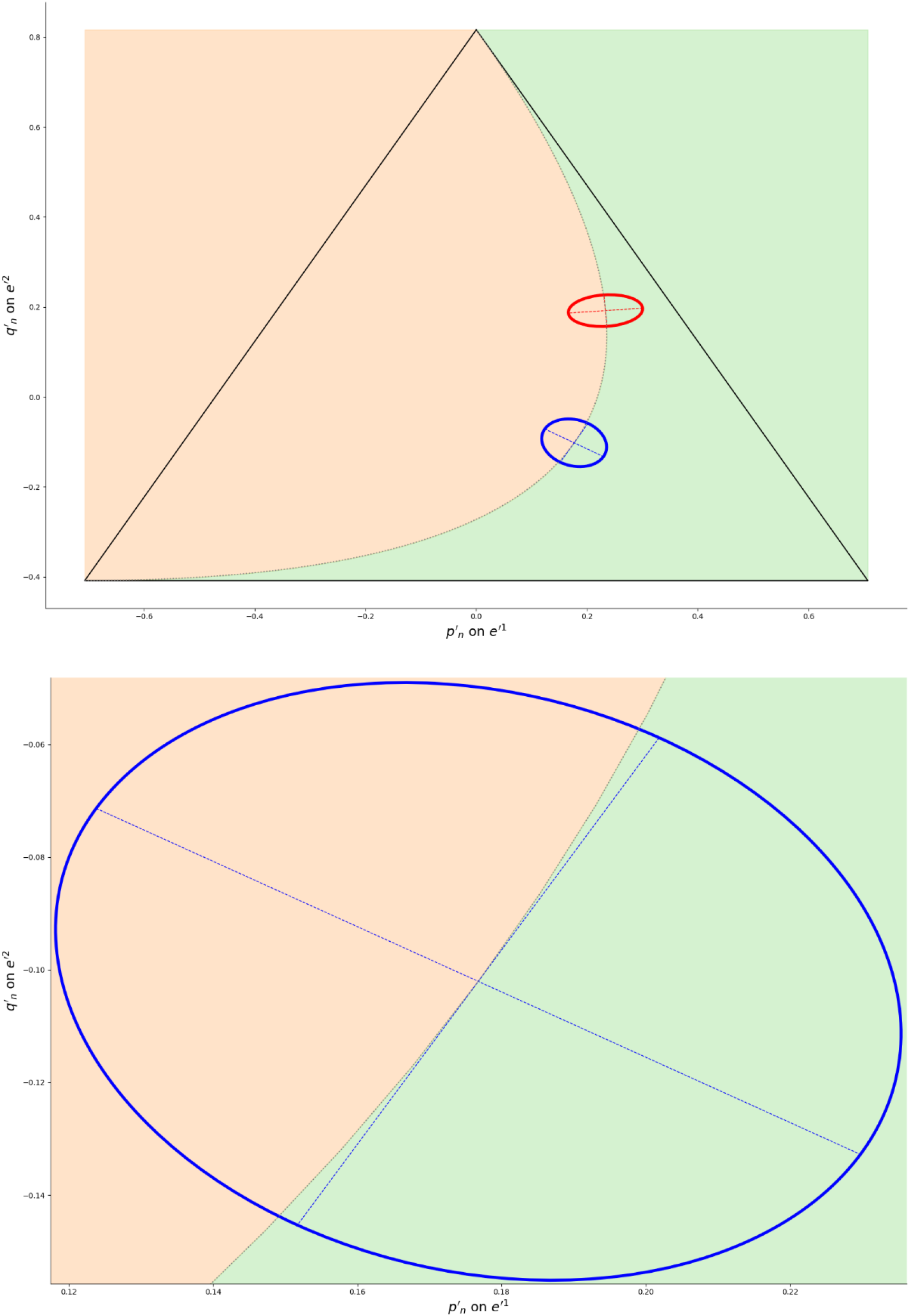
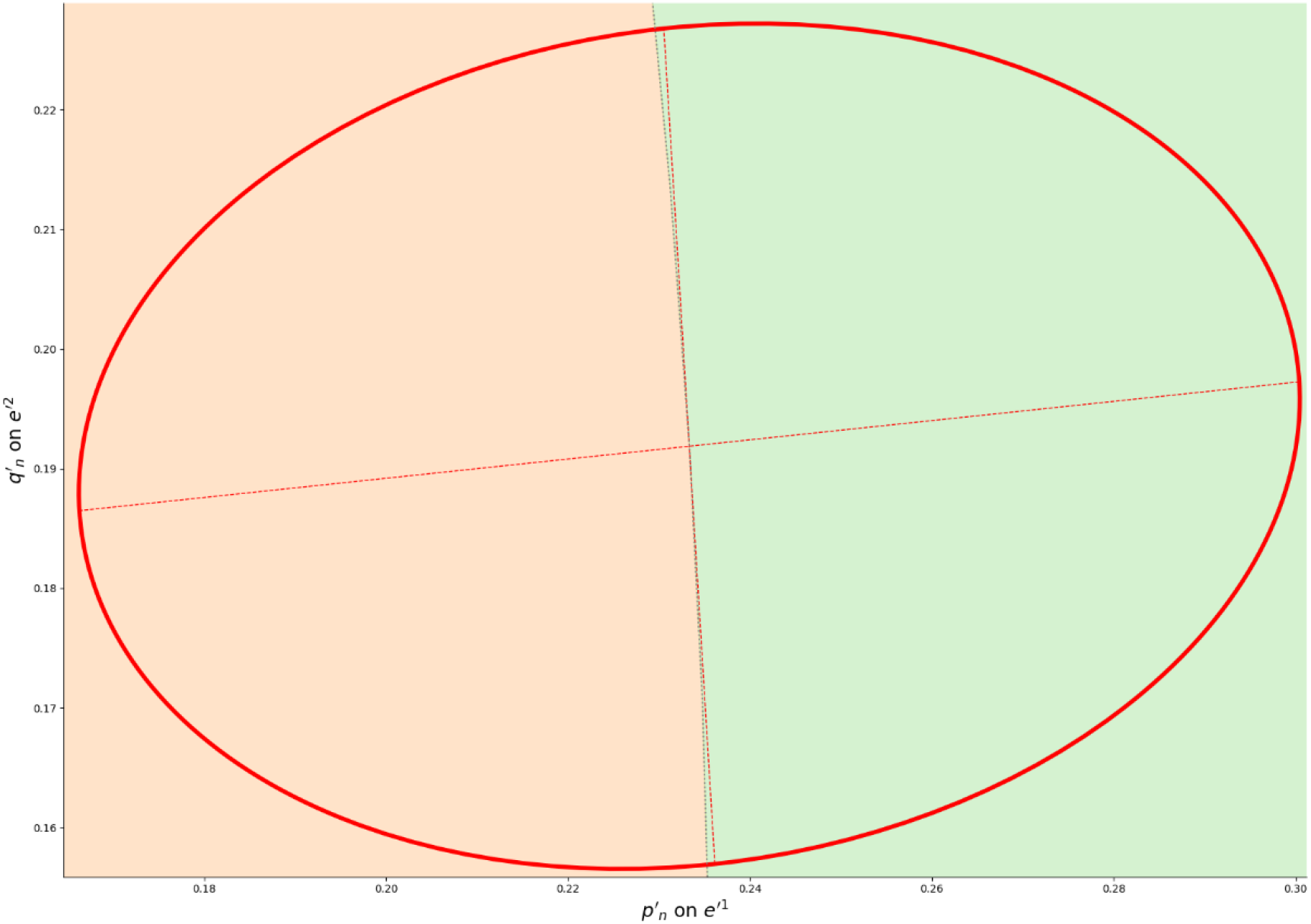
Illustration of why genetic drift tends to increase the probability of negative F_IS_ (area in pastel green) relative to positive F_IS_ (area in pastel orange) due to the concavity of the Hardy-Weinberg parabola (dotted black line; simplex boundaries in solid black) near the equilibrium. Concentration ellipses are shown in color (blue at equilibrium, red on its way to equilibrium), with their two axes as dashed colored lines (equations in Appendix). The top subfigure shows the full simplex in the plane *Π*^*^. The middle and bottom subfigures show close-up view of the concentration ellipses highlighting the relative areas corresponding to negative (pastel green) and positive (pastel orange) F_IS_. The two axes of each concentration ellipse divide its total area into four equal parts. Pastel green (i.e., negative F_IS_) occupies one full half of each ellipse plus a small portion of the opposite half.

